# Resistance, resilience, and functional redundancy of freshwater bacterioplankton communities facing a gradient of agricultural stressors in a mesocosm experiment

**DOI:** 10.1101/2020.04.12.038372

**Authors:** Naíla Barbosa da Costa, Vincent Fugère, Marie-Pier Hébert, Charles C.Y. Xu, Rowan D.H. Barrett, Beatrix E. Beisner, Graham Bell, Viviane Yargeau, Gregor F. Fussmann, Andrew Gonzalez, B. Jesse Shapiro

## Abstract

Agricultural pollution with fertilizers and pesticides is a common disturbance to freshwater biodiversity. Bacterioplankton communities are at the base of aquatic food webs, but their responses to these potentially interacting stressors are rarely explored. To test the extent of resistance and resilience in bacterioplankton communities faced with agricultural stressors, we exposed freshwater mesocosms to single and combined gradients of two commonly used pesticides: the herbicide glyphosate (0-15 mg/L) and the neonicotinoid insecticide imidacloprid (0-60 μg/L), in high or low nutrient backgrounds. Over the 43-day experiment, we tracked variation in bacterial density with flow cytometry, carbon substrate use with Biolog EcoPlates, and taxonomic diversity and composition with environmental 16S rRNA gene amplicon sequencing. We show that only glyphosate (at the highest dose, 15 mg/L), but not imidacloprid, nutrients, or their interactions measurably changed community structure, favoring members of the Proteobacteria including the genus *Agrobacterium*. However, no change in carbon substrate use was detected throughout, suggesting functional redundancy despite taxonomic changes. We further show that communities are resilient at broad, but not fine taxonomic levels: 24 days after glyphosate application the precise amplicon sequence variants do not return, and tend to be replaced by phylogenetically close taxa. We conclude that high doses of glyphosate – but still within commonly acceptable regulatory guidelines – alter freshwater bacterioplankton by favoring a subset of higher taxonomic units (i.e. genus to phylum) that transiently thrive in the presence of glyphosate. Longer-term impacts of glyphosate at finer taxonomic resolution merit further investigation.

## Introduction

Agricultural expansion and intensification are major drivers of global environmental change in both terrestrial and aquatic ecosystems (Song et al., 2018; Springmann et al., 2018; Tilman et al., 2001). Chemicals derived from agricultural landscapes, such as fertilizers and pesticides, are among the main sources of freshwater pollution (Vörösmarty et al., 2010), leading to eutrophication (Carpenter et al., 1998; Keatley, Bennett, Macdonald, Taranu, & Gregory-Eaves, 2011) and biodiversity loss (DeLorenzo, Scott, & Ross, 2001; Relyea, 2009; Stehle & Schulz, 2015). Anthropogenic climate change may intensify these effects as variation in precipitation patterns and increased temperatures affect agrochemicals fate, transport, and behavior in surface and groundwater (Bloomfield, Williams, Gooddy, Cape, & Guha, 2006; Jeppesen et al., 2009). Agricultural runoff to waterbodies particularly increases after storms, acting as a pulse perturbation (Cedergreen & Rasmussen, 2017) while bringing a mixture of nutrients, herbicides and insecticides that may interact to affect aquatic microbial taxa (Flood & Burkholder, 2018) and communities (Lozano & Pratt, 1994; Starr, Bargu, Maiti, & DeLaune, 2017). The impact of agricultural contaminants may depend on whether they are applied alone or in combination (Altenburger, Backhaus, Boedeker, Faust, & Scholze, 2013), and the effects of combinations may be difficult to predict based upon data from single contaminants, possibly due to complex interactions within diverse bacterial communities (Romero, Acuña, & Sabater, 2020).

Agricultural activity has a major impact on bacterioplankton (Kraemer et al., 2020) and, as a consequence, on the ecosystem processes they provide; e.g. decomposition of organic matter (Piggott, Niyogi, Townsend, & Matthaei, 2015) and nutrient cycling (Romero et al., 2020). Altering these processes may have broad consequences for aquatic ecosystem productivity, food webs, and the human activities that depend upon them (Carpenter, Stanley, & Vander Zanden, 2011).

Nutrient pollution is among the most important stressors affecting biodiversity in lakes (Birk et al., 2020). It promotes eutrophication (Smith, Joye, & Howarth, 2006), which can increase bacterial biomass, reduce phytoplankton diversity, and trigger harmful algal blooms (Paerl, Otten, & Kudela, 2018; Smith & Schindler, 2009). While few studies have addressed individual and combined effects of fertilizers with herbicides or insecticides on phytoplankton and zooplankton communities (Baker, Mudge, Thompson, Houlahan, & Kidd, 2016; Chará-Serna, Epele, Morrissey, & Richardson, 2019; Geyer, Smith, & Rettig, 2016), analogous assessments of bacterioplankton are more scarce. Yet, similar to other planktonic communities, bacteria may also be directly or indirectly (e.g. through trophic effects) affected by the individual or combined effects of these stressors, despite not being their intended targets (Muturi, Donthu, Fields, Moise, & Kim, 2017).

The herbicide glyphosate, mainly formulated commercially as Roundup, and the neonicotinoid insecticide imidacloprid (available in different commercial formulations) are among the most commonly used pesticides worldwide (Benbrook, 2016; Simon-Delso et al., 2015), despite restrictions on their use in different jurisdictions. In North America and the European Union, common benchmarks to protect aquatic life range from 800 to 26,600 μg/L of glyphosate for long-term (chronic) exposure, and between 27,000 to 49,000 μg/L for short-term (acute) exposure (CCME, 2012; EFSA, 2016; EPA, 2019). In contrast, lower concentrations of imidacloprid are considered safe for aquatic invertebrates, ranging from 0.009-0.385 μg/L (CCME, 2007; EFSA, 2014; EPA, 2019) (Table S1). Most of these criteria were developed based on toxicity tests on individual eukaryotic organisms, and it remains unclear how bacterial communities respond to these concentrations considered “safe for aquatic life” and what consequences their responses might have on the ecosystem functions they provide.

Glyphosate is a broad-spectrum synthetic phosphonate herbicide used for weed control. It acts by inhibiting the enzyme enolpyruvylshikimate-3-phosphate synthase (EPSPS) involved in the biosynthesis of aromatic amino acids essential to plants, its target group, but also to many fungi and bacteria (Pollegioni, Schonbrunn, & Siehl, 2011). However, some microorganisms are resistant to glyphosate either by expressing an insensitive form of the target enzyme (Funke, Han, Healy-Fried, Fischer, & Schönbrunn, 2006; Healy-Fried, Funke, Priestman, Han, & Scho, 2007) or by metabolizing the molecule and using it as a phosphorus source (Hove-Jensen, Zechel, & Jochimsen, 2014).

Glyphosate could therefore select for resistant species within bacterial communities (Muturi et al., 2017). Moreover, as it may prevent the growth of some phytoplankton species (Smedbol, Lucotte, Labrecque, Lepage, & Juneau, 2017), bacterioplankton could be affected indirectly, for example by reduced competition with phytoplankton.

Unlike glyphosate, imidacloprid is an insecticide commonly used as a seed-coating agent intended to control sapling damage from piercing-sucking insects (CCME, 2007; Jeschke & Nauen, 2008). It acts on insect nervous systems (Roberts & Hutson, 1999) and can be toxic to many aquatic invertebrates, especially insects and crustaceans (Morrissey et al., 2015). Although it is not known to inhibit bacteria directly, it could affect them indirectly via trophic effects on their predators or grazers. If imidacloprid reduces total zooplankton biomass, for example, a reduction in predation pressure could promote an increase in bacterioplankton biomass. Ecosystem functions provided by bacterioplankton, such as carbon use, could subsequently be affected, as has been observed in experiments with other insecticides (Thompson et al., 2016).

Bacterioplankton are important drivers of energy and nutrient cycling in freshwater ecosystems (Falkowski, Fenchel, & Delong, 2008; Konopka, 2009), and more observations are needed to understand how they respond to anthropogenic disturbances (Allison & Martiny, 2008). They may respond with detectable changes in species composition (Allison & Martiny, 2008) that could be permanent, thereby providing a measure of the historical impact of anthropogenic activities on ecosystem health (Kraemer et al., 2020). Alternatively, community composition could be resistant or resilient to changes (Shade et al., 2012). Even if disturbances alter community composition, ecosystem processes may remain stable if pre- and post-disturbance communities are functionally redundant (Allison & Martiny, 2008).

Functional redundancy is thought to be common in microbial communities, as most metabolic pathways controlling biogeochemical cycles are encoded by several different phylogenetic groups. Certain functions, such as photosynthesis and methanogenesis, are however phylogenetically restricted (Falkowski et al., 2008). It is likely that communities are partially redundant for general functions like respiration or biomass production, but non-redundant for more specific functions encoded by unique taxa (Louca et al., 2018). The prevalence of, and reasons for microbial community resistance, resilience, and functional redundancy are still debated (Allison & Martiny, 2008; Shade et al., 2012), particularly in response to novel anthropogenic disturbances which increasingly involve combinations of stressors.

In this study, we experimentally tested the effects of pulse applications of glyphosate and imidacloprid, under low (mesotrophic) or high (eutrophic) nutrient conditions, on bacterioplankton community density, taxonomic composition and richness, and functions related to carbon substrate use. To do so, we filled 1,000 L mesocosms with water and planktonic organisms from a pristine lake located on a mountaintop of a protected area with no history of agricultural activity. Using a regression design, we applied gradients of pesticide concentrations (Fig. 1), spanning ranges observed in surface runoff and freshwater systems (Hénault-Ethier et al., 2017; Morrissey et al., 2015; van Bruggen et al., 2018). Highest doses applied are considered harmful to eukaryotic organisms upon which nationwide water quality guidelines are based (Table S1). To quantify individual and interactive effects of agricultural stressors, we applied these pesticides alone and in combination, and in the presence or absence of nutrient enrichment simulating fertilizer pollution. Pesticides were applied as pulse perturbations to mimic how these contaminants reach natural freshwater ecosystems from agricultural fields, while nutrient enrichments were applied as press treatments to mimic mesotrophic and eutrophic conditions.

**Fig. 1.**
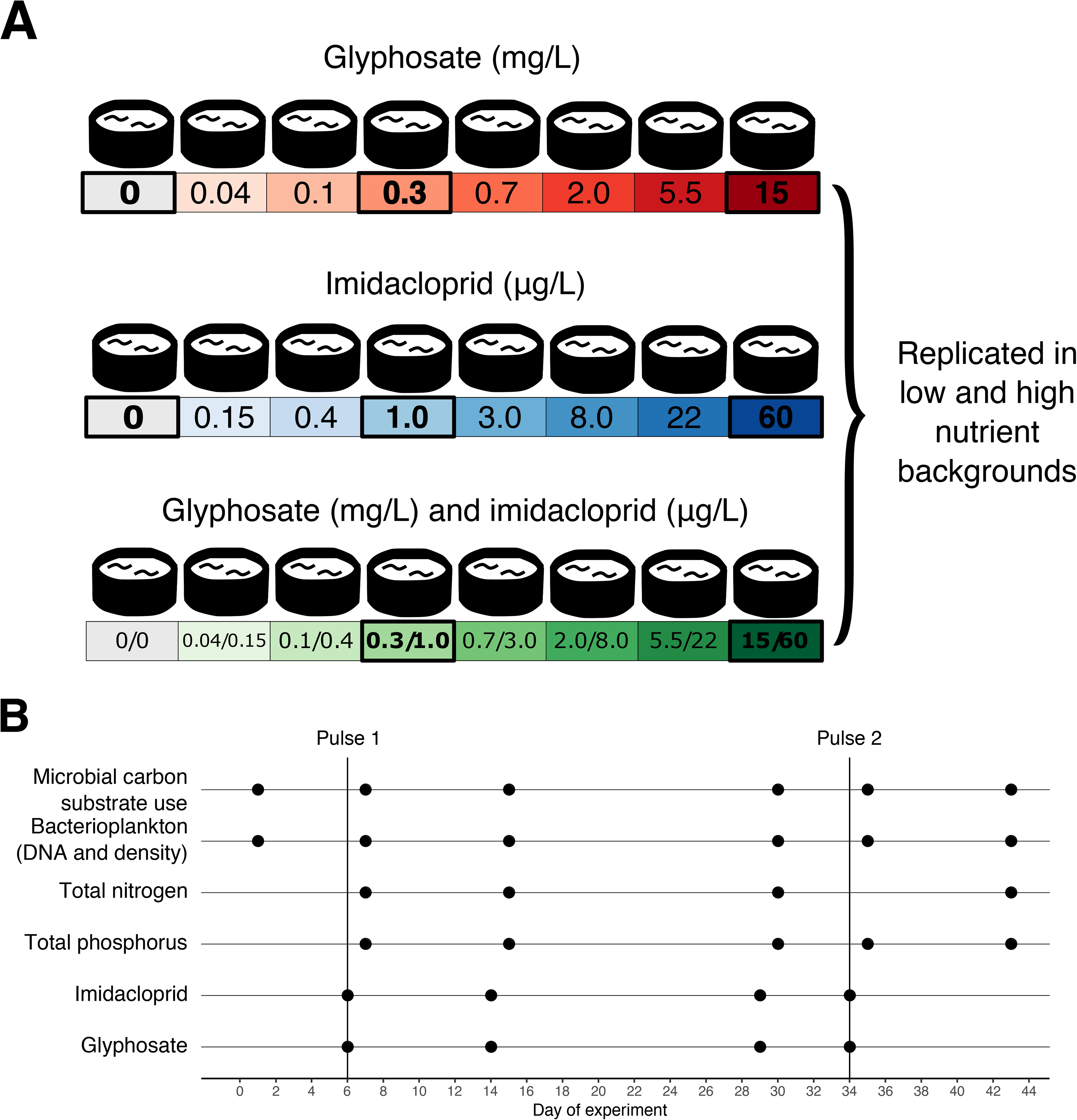
Experimental design and sampling timeline. A) In total, 48 mesocosms (ponds) at the Large Experimental Array of Ponds (LEAP) at the Gault Nature Reserve were filled with 1,000 L of pristine lake water and received two pulses of the pesticides glyphosate and imidacloprid, alone or in combination, at two different nutrient enrichment scenarios. Each box represents an experimental pond and those outlined in bold indicate ponds sampled for DNA extraction and 16S rRNA gene amplicon sequencing. B) The experiment lasted 43 days and pesticides were applied on days 6 (pulse 1) and 34 (pulse 2). Dates of sampling for each variable are indicated with points. Nutrients were added every two weeks at a constant dose, starting seven days before the first sampling day.

We hypothesize that glyphosate will change bacterial community composition and reduce richness, as many taxa depend on the target enzyme (EPSPS) to synthesize aromatic amino acids, while other species encode a resistant allele of EPSPS (Funke et al., 2006; Healy-Fried et al., 2007; Rainio et al., 2021) or are able to metabolize glyphosate (Hove-Jensen et al., 2014). While imidacloprid is less likely to directly impact bacteria, we hypothesize that it can exert indirect effects due to its potential toxicity to aquatic invertebrates (Chará-Serna et al., 2019), releasing grazing pressure on bacterial communities and increasing their density. When applied in combination with glyphosate, imidacloprid may therefore delay or mask the effects of glyphosate on bacterioplankton community structure. Similarly, fertilizers might also increase microbial productivity and mask negative effects of glyphosate, as it does with other contaminants (Alexander, Luis, Culp, Baird, & Cessna, 2013). We also expect some positive effects of glyphosate on bacterial density, as it may serve as a source of phosphorus for some species (Hébert, Fugère, & Gonzalez, 2018; Lu et al., 2020). Finally, we predict that functional diversity will be less prone to changes than taxonomic composition, as bacterial communities tend to be functionally redundant (Louca et al., 2018). We thus expect to detect changes in bacterial community composition and species richness at lower pesticide doses, and changes in functional diversity only at higher doses, or not at all.

## Materials and methods

### Experimental design and sampling

We conducted a mesocosm experiment at the Large Experimental Array of Ponds (LEAP) platform at McGill University’s Gault Nature Reserve (45°32’N, 73°08’W), a protected area with no history of agricultural pollution (Beauséjour, Handa, Lechowicz, Gilbert, & Vellend, 2015) in Quebec, Canada. The pond mesocosms at LEAP are connected to a reservoir that receives water from the upstream Lake Hertel through a 1 km pipe by gravity. On May 11^th^, 2016 (99 days prior to the start of the experiment), 100 ponds were simultaneously filled with 1,000 L of lake water, to acclimate communities to the mesocosm setting. When filling ponds we used a coarse sieve to prevent fish introduction. To maximize initial homogeneity among communities (before treatments), and because this study focuses on planktonic microbes, no sediment substrate was added to the ponds. Tadpoles and large debris such as leaves and pollen were periodically removed with a net before the experiment commenced. Additional lake water was added on a biweekly basis (∼10% of total volume) between May and August to ensure a continuous input of lake bacterioplankton, tracking seasonal changes in the source lake community, and to homogenize communities across ponds. The experiment reported here used 48 of these pond mesocosms from August 17^th^ (day 1) to September 28^th^ (day 43), and it is part of a collaborative experiment that also assessed responses of zooplankton in the same set of ponds (Hébert et al., 2021) and phytoplankton responses in a subset of these ponds for a longer period of time (Fugère et al., 2020).

Throughout our experiment, phosphorus (P) and nitrogen (N) were simultaneously added biweekly to simulate nutrient enrichment at a constant rate, starting on August 10^th^, 7 days before the first sampling day to ensure communities would have passed their exponential growth phase before the first pulse of pesticides was applied. Our nutrient treatment included two levels, with target concentrations of 15 µg P/L (hereafter referred as low-nutrient treatment) typical of mesotrophic Lake Hertel (Thibodeau, Walsh, & Beisner, 2015) and 60 µg P/L (high-nutrient treatment; eutrophic conditions). Nutrient solutions were made using nitrate (KNO_3_) and phosphate (KH_2_PO_4_ and K_2_PO_4_) preserving the same N:P molar ratio (33:1) found in Lake Hertel; the target concentrations were therefore 231 µg N/L and 924 µg N/L for low- and high- nutrient treatments respectively. Over the course of the experiment, the average total P (TP) concentration measured in the source lake was 20.4 µg/L (standard error, SE=±1.3) and the average TP achieved in ponds with no pesticide addition was 13.6 µg/L (SE=±0.71) and 36.7 µg/L (SE=±10.8) respectively for low- and high-nutrient treatment. The average total N (TN) concentration was 556.9 µg/L (SE=±60.7) at Lake Hertel, 407.8 µg/L (SE=±32.7) and 789.0 µg/L (SE=±177.6) respectively in control ponds with low and high nutrient inputs.

Within each nutrient treatment, ponds received varying amounts of the herbicide glyphosate or the insecticide imidacloprid, separately or in combination, in a regression design with seven levels of pesticide concentration plus controls with no pesticide addition (Fig. 1A). The seven levels of target concentration were: 0.04, 0.1, 0.3, 0.7, 2, 5.5 and 15 mg/L for glyphosate and 0.15, 0.4, 1, 3, 8, 22 and 60 µg/L for imidacloprid. There was no replication for each combination of nutrient and pesticide concentration, which is compensated by the wide gradient of pesticides concentration established in the regression design (Fig. 1A). Glyphosate was added in the form of Roundup Super Concentrate (Monsanto©) and target concentrations calculated based on its glyphosate acid content, while imidacloprid was added in the form of a solution prepared with pure imidacloprid powder (Sigma-Aldrich, Oakville, Canada) dissolved in ultrapure water. Treatment ponds received two pulses of pesticides (at days 6 and 34 of the experiment) while nutrients were applied biweekly to maintain a press treatment. The target concentrations of glyphosate and imidacloprid were well correlated with the measured concentrations in the ponds (Fig. S1A-B) with the exception of a few ponds receiving the highest imidacloprid dose which reached lower concentrations than intended, especially after the second pulse (Fig. S1C). These ponds nonetheless reached higher concentrations than ponds lower on the imidacloprid gradients (i.e., a clear gradient was established).

Bacterioplankton communities were sampled at six different timepoints (Fig. 1B): one before pesticide application (day 1); three between pulse 1 and pulse 2 applications (days 7, 15, and 30); and two timepoints after the second pulse (days 35 and 43). Pesticide quantification was performed immediately after each pulse application (days 6 and 34) and at two time points between them (days 14 and 29) while nutrients were quantified on the same days as bacterioplankton except for days one and 35 (Fig. 1B).

Water samples for nutrient and microbial community analyses were collected from each mesocosm with integrated samplers (made of 2.5 cm-wide PVC tubing) and stored in dark clean 1L Nalgene (Thermo Scientific) bottles triple-washed with pond water. To avoid cross contamination, we sampled each pond with a separate sampler and bottle. We kept bottles in coolers while sampling and then moved them to an on-site laboratory, where they were stored at 4 °C until processing, for no longer than 4 hours. Water samples for pesticide quantification were collected immediately after pesticide application (days 6 and 34) in a subset of ponds englobing each gradient and in a smaller subset between the pulses (days 14 and 29). They were stored in clear Nalgene bottles (1 L), acidified to a pH < 3 with sulfuric acid and frozen at -20 °C until analysis.

### Nutrient and pesticide quantification

Quantification of TP and TN from unfiltered water samples were processed at the GRIL (Interuniversity Group in Limnology) analytical laboratory at the Université du Québec à Montréal following standard protocols as outlined by McComb (2002). Duplicate subsamples (40 mL) of water sampled from each pond were stored in acid-washed glass tubes and kept at 4 °C until nutrient concentrations were quantified. TN concentration was determined using the alkaline persulfate digestion method coupled with a cadmium reactor (Patton & Kryskalla, 2003) in a continuous flow analyzer (OI Analytical, College Station, TX, USA). TP was estimated based on optical density in a spectrophotometer (Biocrom Ultrospec 2100pro, Holliston, MA, USA) after persulfate digestion through the molybdenum blue method (Wetzel & Likens, 2000). Glyphosate and imidacloprid concentrations were quantified through liquid chromatography coupled to mass spectrometry using an Accela 600-Orbitrap LTQ XL (LC–HRMS, Thermo Scientific). The method consisted of heated electrospray ionization (HESI) in negative mode for glyphosate, acquisition in full scan mode (50-300 m/z) at high resolution (FTMS = 30,000 m/Dz) and the same LC-HRMS system but using positive HESI mode for imidacloprid (mass range 50-700m/z). Limits of detection were 1.23 and 1.44 μg/L for glyphosate and imidacloprid respectively, while quantification thresholds were respectively 4.06 μg/L, and 4.81 μg/L. Samples falling below limits of detection were pre258 concentrated with a factor of 40X (10 mL samples were reconstituted to 250 μL) and their final concentration were back-calculated according to the concentration factor.

### Estimating bacterial density through flow cytometry

To estimate the density of bacterial cells, we fixed 1 mL of the 1 L sampled pond water with glutaraldehyde (1% final concentration) and flash froze this subsample in liquid nitrogen (Gasol & Del Giorgio, 2000; Ruiz-González et al., 2018). We stored samples at -80 °C until they were processed via a BD Accuri C6 flow cytometer (BD Biosciences, San Jose, CA, USA). After samples were thawed at room temperature (18-20 °C), we prepared dilutions (1:25) with Tris-EDTA buffer (Tris-HCl 10 mM; EDTA 1 mM; pH 8) and aliquoted in two duplicate tubes. Samples were then stained with Syto13 Green-Fluorescent Nucleic Acid Stain (0.1 v/v in DMSO; ThermoFisher S7575) and incubated in the dark at room temperature (18-20 °C) for 10 min. To validate the equipment calibration, we ran BD TruCount Absolute Count Tubes (BD Biosciences) each day, prior to sample processing. Samples were run until reaching 20,000 events, at a rate of 100-1,000 events/s in slow fluidics (14 µL/min). Events within a predefined gate on a 90° light side scatter (SSC-H) versus green fluorescence (FL1-H) cytogram were used for cell counts estimation. This inclusive gate was defined to maximize cell counts accuracy by excluding background noise and large debris. Bacterial density was estimated based on cell counts detected within the gate, flow volume, and sample dilution. We calculated the average bacterial density for each pair of analytical duplicates with a coefficient of variation (CV, i.e., ratio between the standard deviation and average of the duplicate values) less than 0.08. If the CV was greater than or equal to 0.08, the sample was run a third time, and the outlying value was discarded before taking the mean of the two remaining samples.

### Carbon substrate utilization patterns

We used Biolog EcoPlate® assays (Hayward, CA, USA) to infer community-level utilization of dissolved organic carbon by microbes. For all treatments (Fig. 1A) and at each of the six sampled timepoints (Fig. 1B), we added 125 μL of unfiltered pond water to each well of the EcoPlates. Each plate contains, in triplicates, 31 different organic carbon substrates and water controls. These substrates can be grouped into five main guilds (amines/amides, amino acids, carbohydrates, carboxylic acetic acids and polymers), as summarized in Table S2. We measured the optical density at 590 nm in each well as a proxy for microbial carbon substrate use, since it causes a concomitant reduction of the redox-sensitive tetrazolium dye, whose color intensity is measurable at this wavelength. Plates were incubated in the dark at room temperature (18-20 °C) and well absorbance was measured daily until an asymptote was reached (Ruiz-González et al., 2018; Ruiz-González, Niño-García, Lapierre, & del Giorgio, 2015). For each daily measurement, an average well color development (AWCD) was calculated. To correct for variation in inoculum density we selected substrate absorbance values of the plate measurements with AWCD closest to 0.5 (usually after 3-8 days of incubation) as suggested in Garland (2001). We then calculated the blank-corrected median absorbance of each substrate at each sampled timepoint for analyses.

### DNA extraction, 16S rRNA gene amplification and sequencing

We selected a subset of ponds for DNA extraction and subsequent analyses (outlined in bold in Fig. 1A) to assess bacterioplankton community responses at the extremes and the middle of the experimental gradient. From each timepoint and nutrient treatment, we chose two control ponds (beginning of the gradient, no pesticide addition), ponds with the third lowest concentration (middle of the gradient) of each or both pesticides (1 µg/L imidacloprid and/or 0.3 mg/L glyphosate), and ponds with the highest concentration (end of the gradient) used in the experiment for each or both pesticides (60 µg/L imidacloprid and/or 15 mg/L glyphosate). We selected ponds with high concentrations of pesticides to maximize the chance of detecting a response from the bacterial community. That said, we still kept concentrations that fall below available regulatory acceptable concentrations for glyphosate in North America (Table S1), allowing us to ask whether changes in bacterial communities can be detected at concentrations considered safe for aquatic eukaryotes in a region where glyphosate is extensively used (Benbrook, 2016; Simon-Delso et al., 2015). In total, we sampled 16 of the 48 experimental ponds at six timepoints, yielding a total of 96 samples for 16S rRNA amplicon sequencing (Fig. 1B). After sampling 1 L of pond water as described above, we immediately filtered 250 mL through a 0.22 μm pore size Millipore hydrophilic polyethersulfone membrane of 47 mm diameter (Sigma-Aldrich, St. Louis, USA) and stored filters at -80 °C until DNA extraction. We extracted and purified total genomic DNA from frozen filters using the PowerWater DNA Isolation Kit (MoBio Technologies Inc., Vancouver, Canada) following the manufacturer’s protocol, that includes a 5-min vortex agitation of the filter with beads and lysis buffer to enhance cell lysis. We quantified genomic DNA with a Qubit 2.0 fluorometer (ThermoFisher, Waltham, MA, USA) and used 10 ng to prepare amplicon libraries for paired-end sequencing (2 x 250 bp) on two Illumina MiSeq (Illumina, San Diego, CA, USA) runs. We performed a two-step polymerase chain reaction (PCR) targeting the V4 region of the 16S rRNA gene, with primers U515_F and E786_R, as described in Preheim et al. (2013). Further details on PCR reactions, library preparation and amplicon sequencing, including positive controls (mock communities) and negative controls are described in the Supplementary Material.

### Sequence data processing

We used idemp (https://github.com/yhwu/idemp) to demultiplex barcoded fastq files from the sequencing data, and cutadapt to remove remaining Illumina adapters (Martin, 2011). The DADA2 package (Callahan et al., 2016) in R was used to filter and trim reads, using the default filtering parameters with a maximum expected error (maxEE) score of two. Reads were trimmed on the left to remove primers and those shorter than 200 or 150 bp were discarded, respectively, for forward and reverse reads. DADA2 was also used to infer amplicon sequence variants (ASVs), remove chimeras and finally obtain a matrix of ASV counts in each sample for each MiSeq run independently. We used the default parameters of the “learning error rates” function with the multithread option enabled. The number of raw reads and non-chimeric reads obtained from each sample are summarized in Table S3 (average raw reads per sample: 43,159; SE=2,245). Excluding mock communities, extraction blanks and PCR controls, we obtained 1,787,412 raw reads in the first run and 4,702,355 in the second run, of which we retained, respectively, 1,565,021 and 4,188,644 non-chimeric reads. PCR negative controls and extraction blanks produced 214 non-chimeric reads in total; these were excluded from downstream analyses as we only included samples with a minimum of 6,000 reads. Of the 30 expected sequences from the custom mock community (Preheim et al., 2013), DADA2 found 25 exact sequence matches, producing 5 false negatives and 7 false positives (for a total of 32 sequences). In the ATCC mock, 23 of the 24 expected sequences were found, with only 1 false negative but 10 false positives (for a total of 33 sequences). We concatenated DADA2 abundance matrices from each MiSeq run and then used TaxAss (Rohwer, Hamilton, Newton, & McMahon, 2018) to assign ASV taxonomy with a database specifically curated for freshwater bacterioplankton, FreshTrain (Newton, Jones, Eiler, Mcmahon, & Bertilsson, 2011), and GreenGenes (DeSantis et al., 2006), with a minimum bootstrap support of 80% and 50%, respectively. After performing a multiple sequence alignment with the R package DECIPHER (Wright, 2016), we constructed a maximum likelihood phylogenetic tree using the phangorn package following recommendations made by Callahan (2016). For subsequent analyses, we imported the ASV abundance matrix together with taxonomic assignments and environmental data as an object in the phyloseq package (McMurdie & Holmes, 2013) in R. We removed sequence data identified as mitochondria or chloroplast DNA and normalized read counts using the DESeq2 package (Love, Huber, & Anders, 2014), which performs a variance stabilizing transformation without discarding reads or samples (McMurdie & Holmes, 2014), which is important in the context of high read depth variation, as observed among our samples (Table S3). As normalizations such as the DESeq2 method tend to reduce the importance of dominant taxa while inflating the importance of rare taxa (McKnight et al., 2019), for comparison with DESeq2, we additionally normalized the abundance matrix in two ways: (1) by calculating relative abundances (proportions) of each ASV, and (2) by rarefying to 10,000 reads (948 ASVs and 7 samples were consequently removed). These two alternative normalizations are presented in the Supplementary Materials, and are generally concordant with the DESeq2 results in the main text. For most compositional analyses in the main text, we calculated the estimated absolute abundance (EAA) of ASVs per sample by multiplying the DESeq2 normalized ASVs relative abundance by the total bacterial cell counts found in the sample through flow cytometry (Zhang et al., 2017).

### Statistical analyses

To assess resistance and resilience to experimental treatments, we compared changes in bacterial community density, microbial carbon substrate use, as well as bacterioplankton community taxonomic structure (richness and composition), as explained in detail below. We conducted all statistical analyses in R version 3.5.1 (R Core Team, 2008). As we tested hypotheses of different treatment effects at different timepoints, we applied a Bonferroni correction for multiple hypothesis testing.

#### Treatment effects on bacterioplankton density

Time series of bacterial density were analyzed with a generalized additive mixed model (GAMM) with the mgcv R package (Wood, 2017) to quantify the singular and interactive effects of nutrient and of each pesticide treatment on bacterioplankton density as a function of time while accounting for nonlinear relationships. Glyphosate and imidacloprid target concentrations were rescaled (from 0 to 1) to match the scale of the nutrient treatment factor (binary) and we tested for their effect individually or in combination. Individual mesocosms (ponds) were included as a random effect (random smooth) to account for non-independence among measurements from the same pond over time. Model validation was performed by investigating residual distributions, comparing fitted and observed values and checking if basis dimensions (k) of smooth terms were not too low. The model fit (adjusted R^2^) and further details on predictors used in the model, including their statistical significance, are provided in Table S4.

#### Treatment effects on carbon substrate use

We quantified treatment effects on the number of carbon substrates used at each pond and timepoint with a GAMM with the same terms as the GAMM described above for modeling bacterial density. More details are provided in Table S4. To assess the effects of the treatments on carbon substrate utilization patterns by microbial communities over time, we built principal response curves (PRCs) (Auber, Travers-Trolet, Villanueva, & Ernande, 2017). PRCs are a special case of partial redundancy analysis (pRDA) in which time and treatments, expressed as ordered factors, are used as explanatory variables, while community composition is the multivariate response. Time is considered as a covariable (or conditioning variable) whose effect is partialled out, and changes in community composition with the treatments over time are always expressed as deviations from the control pond at each timepoint. PRCs also assess the contribution of each species to the treatment effect through the taxa weight (also known as species score) usually displayed in the right y-axis of a PRC diagram (Van den Brink, den Besten, bij de Vaate, & ter Braak, 2009). The significance of the first PRC axis was inferred by permuting the treatment label of each pond while keeping the temporal order, using the permute R package (Simpson, 2019) followed by a permutation test with the vegan R package (Oksanen et al., 2018). Before performing PRCs we transformed the community matrix (containing carbon substrate use data) using the Hellinger transformation (Legendre & Gallagher, 2001). PRC of community carbon utilization patterns was performed for the 31 substrates individually and grouped into five guilds (Table S2).

#### Treatment effects on bacterioplankton community taxonomic structure

To infer the impact of treatments on bacterioplankton taxonomic diversity over time, we calculated alpha diversity as richness (number of observed ASVs) and as the exponent of the Shannon index (or Hill numbers (Jost, 2006)) of each sample after rarefying the ASV abundance matrix to 10,000 reads without replacement and modelled their response to pesticide and nutrient treatments using GAMMs. Model equations, their fit (adjusted R^2^) and statistics of significant terms are reported in Table S5. In this analysis, pesticides treatments were considered factors (low vs. high) because 16S rRNA reads data were only available for a subset of concentrations (Fig. 1A). Pesticides and nutrient treatments were coded as ordered factors and models were validated after investigation of residual distributions, comparison of fitted and observed values and checking if the basis dimension (k) of smooth terms was sufficiently large.

To assess differences in community composition, we calculated weighted UniFrac distances (Lozupone & Knight, 2005) and Jensen-Shannon divergence (JSD) among the subset of samples selected for DNA analyses and represented them in principal coordinate analysis (PCoA) bidimensional plots. These two metrics are complementary as the first is weighted for phylogenetic branch lengths unique to a particular treatment, and the second assesses changes in community composition at the finest possible resolution, tracking ASVs regardless of their phylogenetic relatedness. We performed a series of permutational analyses of variance (PERMANOVA) based on weighted UniFrac distances and JSD to test the effect of treatments (as factors) on community composition at four sampled timepoints separately: at day 1 (before any treatment was applied), day 7 (immediately after the first pulse), day 15 (11 days after the first pulse), day 30 (immediately before the second pulse) and day 43 (last day of the experiment, after the second pulse). We also performed an analysis of multivariate homogeneity (PERMDISP) to test for homoscedasticity in groups dispersions (Anderson, 2006) because the PERMANOVA may be sensitive to non-homogeneous dispersions within groups and thus mistake it as among-group variation (Anderson, 2001). A significant PERMDISP (*p*<0.05) indicates different within-group dispersions and thus should be used in combination with visual inspection of the ordination plots to interpret the PERMANOVA results.

Using EAA after read depth normalization with DESeq2, we further visualized bacterioplankton community temporal shifts with PRCs, asking if the extent of community turnover varied across phylogenetic levels. Separate models were built for ASVs grouped at various phylogenetic levels, from phylum to genus. For each PRC model, we evaluated the proportion of variance (inertia) explained by the conditional variable (time) and the constrained variable (treatments), as well as the proportion of explained variance per axis (the eigenvalue of each RDA axis divided by the sum of all eigenvalues). We used these values to decide which PRC model, if at the phylum, class, order, family, genus or ASV level, best explained the variation in the data, and we tested for the significance of the first PRC axis through a permutation test with the permute and vegan packages in R (Oksanen et al., 2018; Simpson, 2019). Taxa weights representing the affinity of the most responsive taxa with the treatment response curve are displayed the right y-axis of each PRC diagram. Before performing each PRC we transformed the community matrix using the Hellinger transformation (Legendre & Gallagher, 2001). The abundance of the three genera with the highest PRC taxa weights were modeled with GAMMs to explore how treatments impacted their (potentially non-linear) abundances over time, and to provide further validation of the treatment effects detected by PRCs. The GAMM response variable was the log-transformed (log (1+x), where x is the variable) EAA of each of the three genera, after reads had been rarefied to 10,000 reads per sample without replacement. We opted for using rarefied data instead of DESeq2 normalization which is intended for community analyses (Weiss et al., 2017) and the GAMMs focused on specific taxa of interest. Modeled abundances were visualized with the R package itsadug (Van Rij, Wieling, Baayen, & van Rijn, 2020).

## Results

### Bacterial cell density is weakly affected by glyphosate while microbial community carbon substrate utilization is resistant to all stressors

Overall bacterial cell density showed a strong but non-linear increase over time across all ponds (GAMM, effect of time: F=17.5, *p*<0.001, Table S4; Fig. 2). The time-independent effect of nutrients on bacterial cell density was weak but positive (GAMM, t=4.1, *p*<0.001), and, over time, glyphosate had a weak positive effect on bacterial density (GAMM, factor-smooth interaction between time and glyphosate: F=6.6, *p*<0.001) (Table S4). The interactive effect of nutrients and glyphosate was also weak, and not significant after Bonferroni correction for multiple testing (GAMM, F=5.7, uncorrected *p*=0.018, Table S4). Overall, these results indicate that, despite increasing over time across ponds, bacterioplankton densities also slightly increased in response to nutrient and glyphosate addition.

**Fig. 2.**
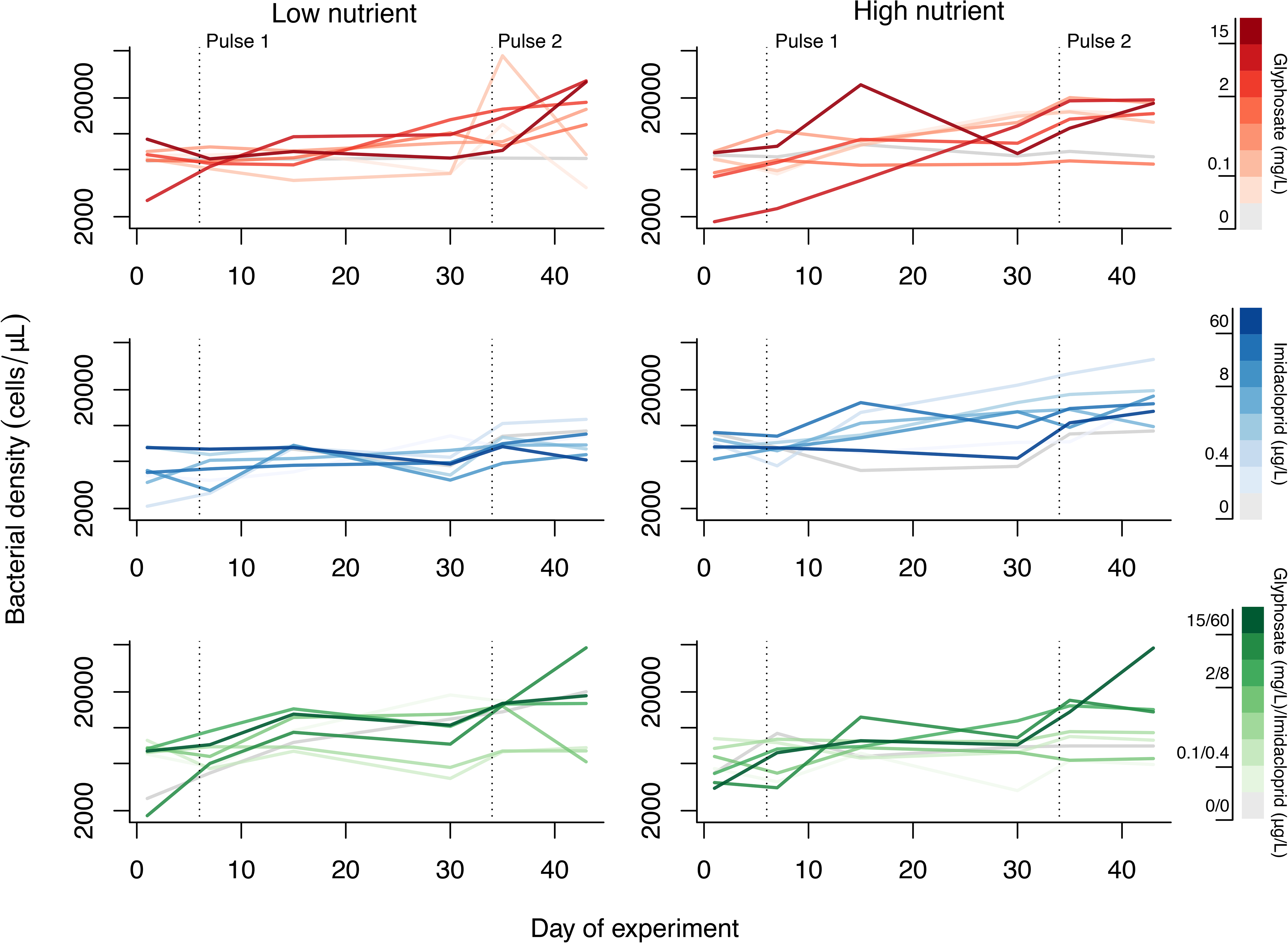
Bacterial cell density dynamics during the experiment. Total bacterial density is plotted over time in ponds with low- or high- nutrient enrichment. Dashed vertical lines indicate days of pesticide pulses application. Ponds with both glyphosate and imidacloprid follow the same gradient pattern as treatments with either of these pesticides applied alone.

The number of carbon substrates used by the microbial community diminished slightly over time (GAMM, F=6.0, *p*<0.001, Table S4). However, neither glyphosate, imidacloprid, nutrients, nor their interactions had significant effects on carbon substrate utilization as assayed by EcoPlates (Table S4). In addition, the PRC analysis did not reveal any significant treatment effects on microbial utilization of any of the 31 unique carbon substrates when considered separately (Fig. 3A; permutation test for the first constrained eigenvalue, F=12.28 *p*=0.295) or when grouped into guilds (Fig. 3B; F=34.46 *p*=0.355). To simplify visualization and facilitate comparison with treatments selected for community taxonomic characterization, the PRCs in Fig. 3 included the same ponds as those used for DNA analyses. PRCs including all ponds in the tested gradient showed similar results (Fig. S2A and Fig. S2B). We conclude that, despite slight changes in the number of substrates being used over time, none of the treatments significantly affected microbial community-level carbon utilization profiles.

**Fig. 3.**
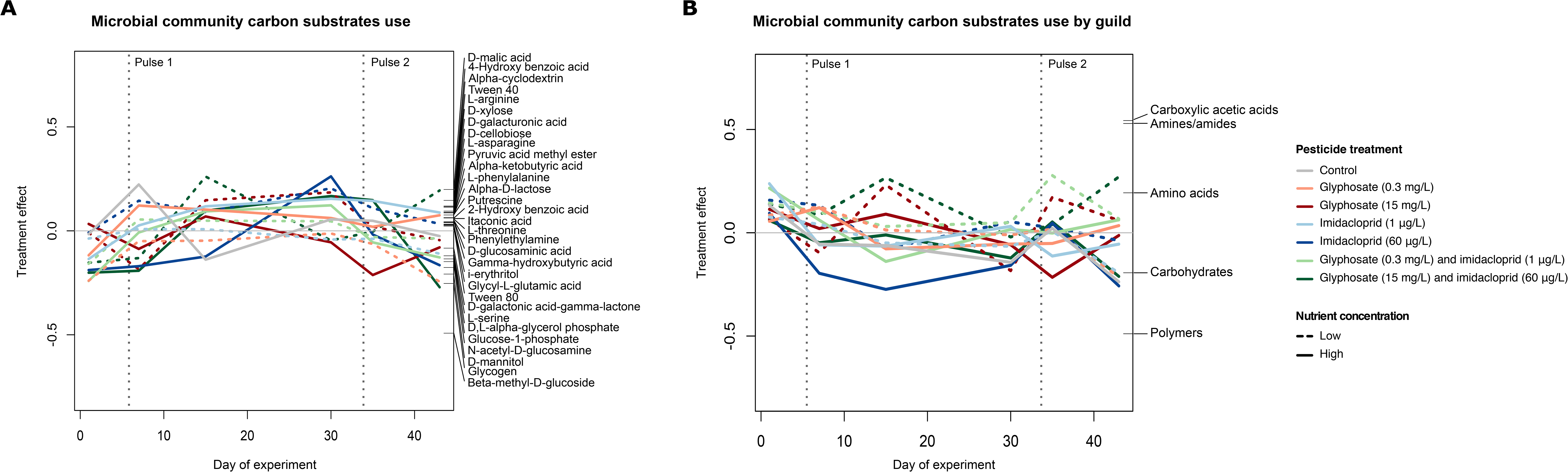
Microbial community carbon substrate utilization. Principal response curves (PRCs) of selected experimental treatments show no significant difference between controls and pesticide treatments when microbial communities are described according to (A) their ability to metabolize 31 different carbon substrates when analysed individually or (B) when grouped into guilds. Weights of each tested compound or guild are shown along the Y axis (right). Dashed vertical lines indicate days of pesticides pulses application. For ease of comparison, the PRCs were calculated based on the subset of samples for which DNA was extracted. The PRC displayed in (A) explains 15.1% of the variation while the one displayed in (B) explains 42.2%, suggesting that grouping substrates into guilds improves the explanatory power of the PRC.

### Bacterioplankton community structure responses

#### Glyphosate has a minor time-independent effect on community diversity and a ***major effect on community composition over time***

We calculated two metrics of bacterioplankton community alpha diversity in each sample: taxon richness, estimated as the logarithm of the total number of observed ASVs after rarefying (Fig. 4A), and the exponent of the Shannon index, which combines information about ASV richness and evenness (Fig. 4B). No significant time-dependent effect of any treatment was detected, although ponds with high glyphosate concentration (15 mg/L) had a lower Shannon diversity when averaged across all timepoints (GAMM, t=-3.51, *p*=0.001, Table S5), and the same was observed for ASV richness but with a non-significant effect after multiple test correction (GAMM, t=-2.89, uncorrected *p*=0.006, Table S5). Overall, the effect of glyphosate on bacterioplankton alpha diversity was relatively weak and not influenced by time (Table S5).

**Fig. 4.**
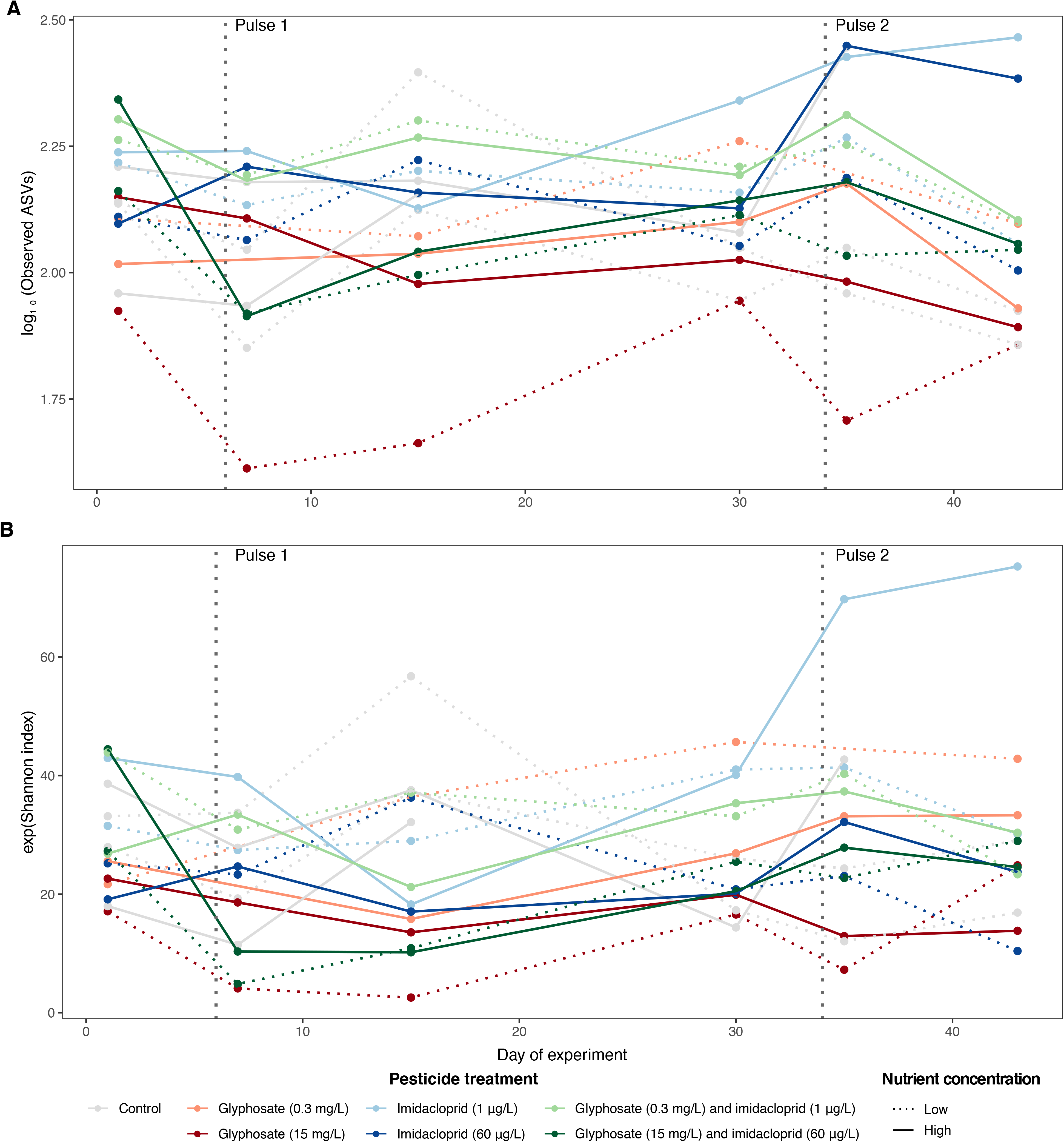
**Bacterioplankton alpha diversity variation across experimental treatments over time**, calculated as (A) the (log transformed) observed number of ASVs per sample and as (B) the exponent of Shannon index. Dashed vertical lines indicate days of pesticides pulses application.

We also tracked changes in bacterioplankton community composition, in two ways: with weighted UniFrac distance and JSD, both calculated after normalizing read depth per sample with DESeq2 (or with alternative normalizations described below). We display these changes in community composition using PCoA, with a separate plot for each timepoint of the experiment (Fig. 5). Glyphosate explained a significant proportion of the variation in both metrics of community composition, with R^2^ ranging from 0.29 to 0.58, depending on the time following glyphosate application (PERMANOVA, *p*<0.007 for both metrics at all tested timepoints after pesticide pulses, except for weighted UniFrac distance at day 30, Tables S6 and S7). Nutrients and imidacloprid did not significantly affect community composition, alone or in combination with other treatments (Tables S6 and S7). Although nutrients appear to have a slight effect on community composition on day 15 (uncorrected *p*=0.027 for weighted UniFrac and JSD, Table S6 and Table S7) and on day 30 (uncorrected *p*=0.055 for weighted UniFrac, Table S6, and uncorrected *p*=0.013 for JSD, Table S7), the effect is not significant after Bonferroni correction, and the explained variance is never as high as it is for glyphosate on the same day (R^2^=0.12 for both weighted UniFrac and for JSD at both days, Table S6 and Table S7). We conclude that glyphosate was the dominant driver of compositional changes as it produced a significant and consistent effect on bacterioplankton communities, independent of other stressors, on days 7, 15, 30 and 43 according to JSD, and on days 7, 15 and 43 according to weighted UniFrac distance.

**Fig. 5.**
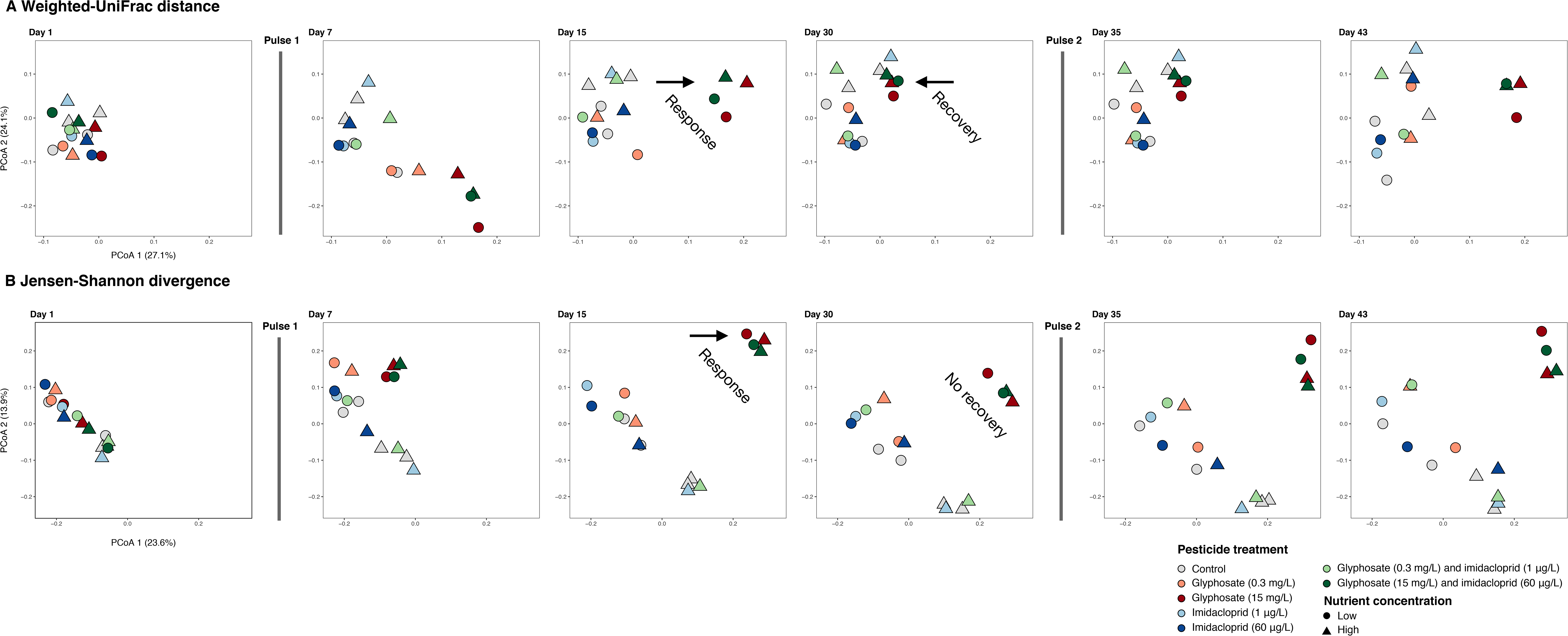
**PCoA ordinations of bacterioplankton community composition in response to experimental treatments**, based on (A) weighted UniFrac distance or (B) Jensen-Shannon divergence calculated on ASV estimated absolute abundances after a DESeq2 normalization. Dashed vertical lines indicate days of pesticides pulses application. Each sampling day is plotted in a separate panel to facilitate visualization of treatment effects on community composition, mainly driven by high glyphosate (15 mg/L).

Alternative read depth normalization methods (ASV relative abundance and rarefied data; see Methods) produced qualitatively similar results, showing the predominant effect of glyphosate on community composition (Fig. S3), with a slight delay in the effect of the first glyphosate pulse compared to the DESeq2 normalization (Fig. 5). The effect of glyphosate on bacterioplankton community composition is detected regardless of the data normalization (Table S8), but is more apparent using DESeq2 (compare Fig. 5 to Fig. S3). This might be because DESeq2 involves a log transformation which reduces the weight of highly abundant community members (McKnight et al., 2019). If less abundant taxa are more responsive to glyphosate, this could explain why this effect is more apparent with DESeq2 normalization.

#### Bacterioplankton communities recover over time at broad phylogenetic scales *from the first glyphosate pulse*

On day 30 (24 days after the first pesticide pulse and before the second pulse), the bacterioplankton community composition in ponds that had been affected (on day 15) by a high dose of glyphosate (15 mg/L) appeared to recover according to weighted UniFrac (Fig. 5A), but not when using JSD applied to ASVs (Fig. 5B). Using weighted UniFrac, the effect of glyphosate was visibly weaker on day 30 (Fig. 5A) and at the limit of significance after Bonferroni correction (PERMANOVA, R^2^=0.29, uncorrected *p*=0.007, Table S6), but still significant using JSD (PERMANOVA, R^2^=0.34, *p*=0.001, Table S7). Viewed together, our series of ordinations show that detection of community recovery depends upon whether phylogenetic information is taken into account. Recovery was apparent when phylogenetic distance among ASVs was calculated (as measured by UniFrac distance, on day 30, control and high-glyphosate communities approach each other, Fig. 5A) but undetected at the ASV level, independent of phylogeny (as measured by JSD, differences between control and high-glyphosate communities keep significant on day 30, Fig. 5B). As such, the community appears to be resilient at a broad phylogenetic level, but not at the finer ASV level, indicating that glyphosate-sensitive ASVs are replaced with phylogenetically-close relatives.

To further assess how resilience varied at different phylogenetic scales, we used PRCs to track community changes at the phylum and ASV levels (Fig. 6). Given that nutrient inputs were not major drivers of community composition (Tables S6 and S7), we built PRCs by combining ponds with the same pesticide treatment, irrespective of nutrient load. This facilitated the visualization of pesticide effects, while capturing the same effects as PRCs considering all experimental treatments separately (compare Fig. 6A and Fig. S4). We further compared PRCs at different phylogenetic scales, from class to genus level (Fig. S5). PRCs captured a significant amount of the variation in community responses to pesticide treatments over time (phylum level: F=31.22, class: F=34.28, order: F=26.19, family: F=21.30, genus: F=20.6, ASV: F=10.61, all *p*=0.001; Table S9), with greater variation explained at broader taxonomic levels compared to finer levels. The variance explained by the first PRC axis decreased from 47.7% at the phylum level to 22.1% at the ASV level (Table S9). At the broadest taxonomic scale (phylum), communities showed a clear response to high (15 mg/L) but not low (0.3 mg/L) concentrations of glyphosate, followed by a recovery before the second pulse (Fig. 6A). Notably, no recovery was observed at the ASV level (Fig. 6B), consistent with the community composition analysis (Fig. 5). Imidacloprid had no detectable effect at any concentration, whereas the highest concentration of glyphosate caused the greatest effect on bacterioplankton communities. Similar response and recovery patterns were also observed down to the genus level, with progressively weaker recovery at finer taxonomic scales (Fig. S5). Community composition showed recovery 24 days after the first pulse of glyphosate, but failed to recover after the second pulse (Fig. 5A and Fig. 6A). While this does not exclude the possibility of an eventual recovery, the duration of our experiment (which ended nine days after the second pulse) was likely insufficient to permit subsequent recovery. These results further support that high concentrations of glyphosate led to long-lasting community shifts at the ASV or genus level, whereas community resilience can be achieved at broader phylogenetic scales.

**Fig. 6.**
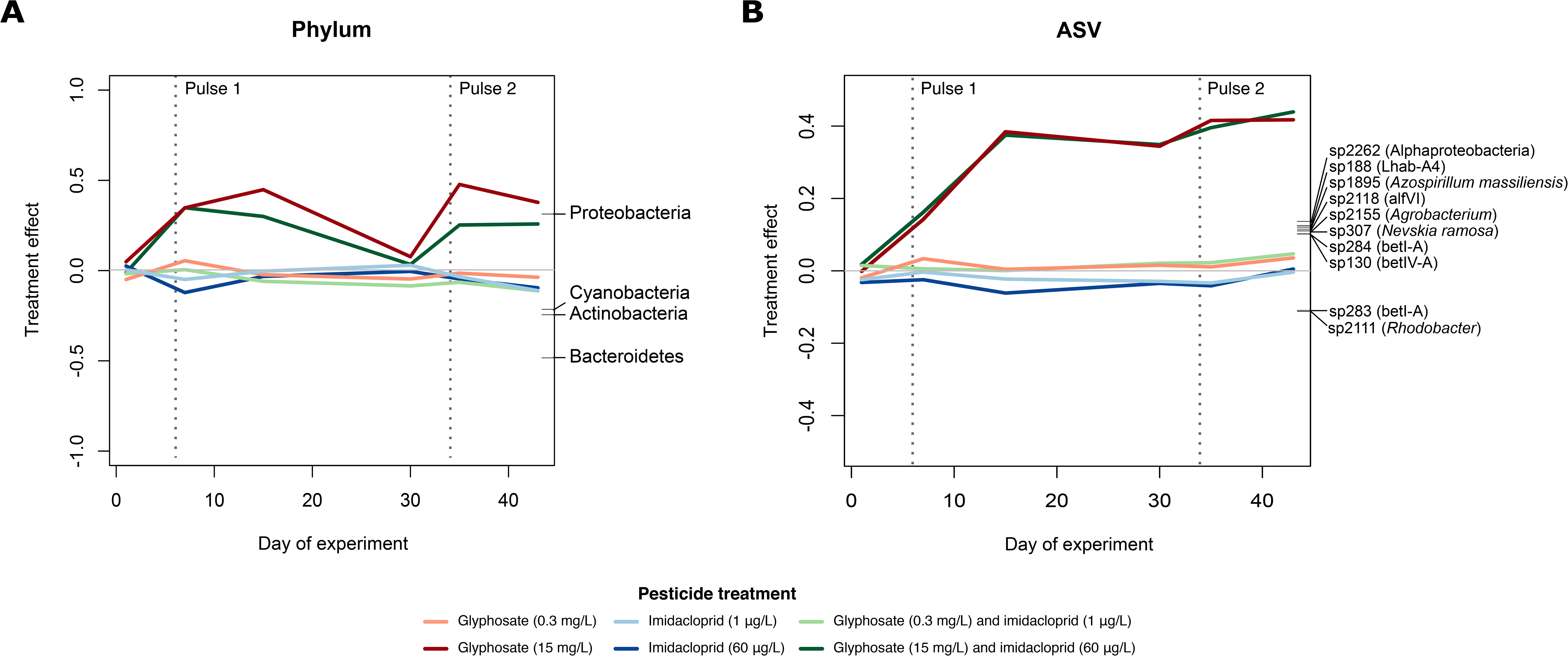
PRCs showing the effect of pesticide treatments over time relative to control ponds at (A) the phylum level or (B) the ASV level. Dashed vertical lines indicate days of pesticides pulses application. Only phyla with weights >0.2 and ASVs with weight >0.1 are plotted on the right Y axis to facilitate visualization. The finest available annotated level of taxonomic assignment of each ASV is indicated in panel B. Low- and high-nutrient treatments were grouped together for clarity, but follow the same pattern when considered separately (Fig. S4). See Fig. S5 for PRCs at other taxonomic levels. These analyses were based on ASV estimated absolute abundances after a DESeq2 normalization.

#### Dynamics of the taxa most responsive to treatments

The phylum Proteobacteria was the most positively affected by glyphosate (Fig 6A; Table S10), with relative abundance over 60% in the high glyphosate treatment (15 mg/L) and ∼50% or less in other treatments and controls (Table S11). Bacteroidetes, Actinobacteria and Cyanobacteria were the most negatively affected phyla (Fig. 6A; Table S10, Table S11). Of the ten ASVs with the highest absolute taxa weights, all belonged to the phylum Proteobacteria (Fig. 6B, Table S12) and, except for sp283 and sp2111, they were all positively affected by glyphosate. An ASV assigned to the genus *Agrobacterium* was among the ASVs that responded most positively to high glyphosate treatment (Fig. 6B; Table S12). The GAMM showed that ASVs assigned to the genus *Agrobacterium* increased in EAA over time in ponds receiving high doses of glyphosate (GAMM, factor-smooth interaction between time and high glyphosate treatment: F=19.49, *p*<0.001, Table S13), or receiving both high glyphosate and imidacloprid (GAMM, factor-smooth interaction between time and treatment with high concentrations of both glyphosate and imidacloprid: F=20.66, *p*<0.001, Table S13). A linear time-independent effect of glyphosate was also detected in experimental ponds treated with the highest concentrations of both pesticides together (GAMM, t=7.50, *p*<0.001, Table S13) or glyphosate alone (GAMM, t=6.25, *p*<0.001, Table S13). The modeled *Agrobacterium* abundance (Fig. S6A) shows a similar ’response followed by recovery’ pattern over time as the overall community response at the phylum level (Fig. 6A), suggesting that the positive effect of glyphosate on Proteobacteria may be driven by *Agrobacterium*.

The other two most positively affected genera (*Flavobacterium* and *Azospirillum*, Fig. S5D) increased in abundance in response to the combination of glyphosate at 15 mg/L and imidacloprid at 60 μg/L (GAMM, factor-smooth interaction between time and treatment with high concentrations of both glyphosate and imidacloprid on *Flavobacterium*: F=17.35, *p*<0.001, and on *Azospirillum*: F=6.27 p=0.001, Table S13) or glyphosate alone at 15 mg/L (GAMM, factor-smooth interaction between time and high glyphosate treatment on *Flavobacterium*: F=3.63, *p*=0.031, not significant after Bonferroni correction; and on *Azospirillum*: F=5.41, *p*=0.002, Table S13), but the effects were not as strong as detected for *Agrobacterium* (Table S13). In contrast to the recovery pattern observed in *Agrobacterium* exposed to both the independent and combined highest concentrations of glyphosate (Fig. S6A), the modeled abundance of *Flavobacterium* (Fig. S6B) and *Azospirillum* (Fig. S6C) followed distinct patterns in these two treatments. *Flavobacterium* responded weakly to high doses of both pesticides, mainly after the second pulse, whereas *Azospirillum* recovered partially after responding to the first pulse, but only in ponds treated with the highest concentrations of both pesticides. Despite the overall strong effect of glyphosate on Proteobacteria, these results highlight how different bacterioplankton taxa (including *Agrobacterium* and *Azospirillum –* both Alphaproteobacteria) can show subtly different responses and recovery patterns to pesticides.

## Discussion

### Context and summary of the experiment

The herbicide glyphosate has been shown to affect aquatic microbial community structure in a variety of natural environments and experimental setups (Berman et al., 2020; Lu et al., 2020; Muturi et al., 2017; Stachowski-Haberkorn et al., 2008). Likewise, the insecticide imidacloprid may disrupt aquatic food webs (Yamamuro et al., 2019), with potential, yet poorly explored consequences for bacterioplankton. The interactive effects of these pesticides on bacterioplankton – and how they might vary depending on fertilizer use and lake trophic status – are relevant because such agrochemical mixtures are common in agriculturally impacted watersheds. Here, we tested how individual and combined gradients of glyphosate and imidacloprid affected bacterioplankton communities in aquatic mesocosms receiving different nutrient inputs. Although they are incomplete representations of natural ecosystems, mesocosm experiments allow us to manipulate and replicate the exposure of complex lake bacterial communities to agricultural chemical pollutants commonly found in freshwaters (Alexander, Luiker, Finley, & Culp, 2016). The current experiment is limited to the response of bacterioplankton communities derived from a pristine lake. Future studies focusing on biofilms and sediments could complement our results, as many contaminants accumulate in lake sediments and may affect the biofilm community structure (Fernandes et al., 2019; Khadra, Planas, Girard, & Amyot, 2018; Romero et al., 2020).

### Glyphosate as a driver of community structure

Our data support the prediction that glyphosate would affect bacterioplankton community structure, which occurred at the highest tested concentration (15 mg/L). Contrary to expectation, no evident interaction between glyphosate and imidacloprid or nutrient load was detected in determining either bacterial density or community structure. High doses of glyphosate resulted in a weak time-independent reduction of bacterioplankton alpha diversity, and a more pronounced change in community composition over time. As hypothesized, glyphosate and nutrient treatments slightly increased bacterial density, suggesting a mild fertilizing effect of glyphosate consistent with it being a potential phosphorus source (Hove-Jensen et al., 2014; Lu et al., 2020). Most bacterioplankton from a pristine source environment (Lake Hertel) are thus able to cope with concentrations of imidacloprid as high as 60 μg/L and of glyphosate as high as 0.3 mg/L, but they may be sensitive to glyphosate concentrations exceeding 15 mg/L. The regulatory criteria intended for eukaryotes (below 60 μg/L for imidacloprid; Table S1) were sufficient to preserve bacterioplankton diversity in the experimental conditions at LEAP. On the other hand, the threshold of 15 mg/L for glyphosate deserves further attention from regulatory agencies, as this concentration impacted bacterioplankton composition, which is known to affect lake health and freshwater quality (Kraemer et al., 2020).

Although the highest targeted imidacloprid concentration was not always achieved in all ponds (Fig. S1), this cannot entirely explain its lack of detectable effect on bacterioplankton. Community composition of ponds receiving measured concentrations of imidacloprid as high as 15 µg/L or more did not deviate from controls, confirming a true lack of effect at least up to that concentration. Alternatively, the absence of a detectable response might be due in part to rapid degradation of imidacloprid in water, which fell below the limit of detection between pulses (Fig. S1). The absence of a bacterioplankton response is also consistent with the weak or undetectable response of zooplankton biomass to imidacloprid pulses in the same experiment (Hébert et al., 2021). The invertebrate community in the experimental ponds was mainly composed of the zooplanktonic groups Cladocera, Copepoda and Rotifera, and only copepods declined over time after pulse 2, with no resulting effect in total zooplankton biomass (Hébert et al., 2021). Overall, these results indicate that the concentrations of imidacloprid applied in this experiment were not sufficient to strongly alter either zooplankton or bacterioplankton biomass or community structure.

Our results suggest that two properties of ecological stability – resistance and resilience – are at play in lake bacterioplankton: functions related to microbial carbon substrate use are resistant to imidacloprid, glyphosate and their interactions in different nutrient backgrounds, while bacterioplankton community composition is resilient following disturbance caused by a glyphosate pulse at 15 mg/L. The recovery of bacterioplankton community composition was only evident when grouping ASVs at higher (more inclusive) taxonomic or phylogenetic levels. Glyphosate thus drove a turnover of bacterioplankton ASVs which, even after the recovery, are different from the ASVs initially found in the undisturbed community.

### Proteobacteria are major responders to glyphosate

Glyphosate treatments had a strong positive effect on the phylum Proteobacteria, previously found to be favoured by high concentrations of glyphosate in rhizosphere- (Newman et al., 2016) and phytoplankton-associated communities (Wang, Lin, Li, Lin, & Lin, 2017). Multiple species of Proteobacteria can use glyphosate as a source of phosphorus by breaking its C-P bond (Hove-Jensen et al., 2014). We identified *Agrobacterium*, a genus of *Rhizobiaceae* containing species known to degrade glyphosate (Hove-Jensen et al., 2014), as being highly favored in the glyphosate treatment at 15 mg/L. The abundance of ASVs assigned to this genus peaked after each pulse and decreased before the second pulse, coinciding with the community recovery observed 24 days after the first perturbation. The ability to degrade glyphosate may be widespread in the family *Rhizobiaceae* (Liu, McLean, Sookdeo, & Cannon, 1991), and *Agrobacterium* have also been found to encode glyphosate-resistant EPSPS genes (Funke et al., 2006). In fact, this genus was used to create glyphosate-resistant crops, i.e. the so-called ‘Roundup-ready technology’ (Funke et al., 2006). While glyphosate may be a stressor for the microbial community at large (e.g., phytoplankton (Fugère et al., 2020)), it may be a resource for some members such as *Agrobacterium*, who could potentially detoxify the environment and thus facilitate community recovery after a pulse perturbation. Further genomic and metagenomic analyses of our experimental samples could reveal whether these ecological dynamics are underlain by evolutionary adaptation, and whether community resistance and resilience can be explained by the initial presence of resistant bacteria in the community, or to *de novo* mutations or gene transfer events.

Glyphosate could have driven changes in the bacterial community via direct mechanisms (e.g. by affecting species with a sensitive EPSPS, its target enzyme) or indirect mechanisms (e.g. effects on other trophic levels that cascaded down to bacteria via predation or other interactions). In a previous study of the same experiment described here that focused on the responses of eukaryotic phytoplankton, we found that glyphosate treatment reduced the diversity of phytoplankton, but did not significantly change phytoplankton community composition (Fugère et al., 2020). Although a reduced phytoplankton diversity could indirectly affect bacterioplankton community composition, a direct effect of glyphosate on bacteria seems more plausible as the taxa favored by the treatment (mainly Proteobacteria) have been previously shown to be directly affected in a similar way (Janßen et al., 2019; Wang et al., 2017). Indeed, bacterial degradation of glyphosate likely released bioavailable phosphorus, stimulating phytoplankton growth (Fugère et al., 2020). Further studies will be needed to disentangle how the effects of pesticides cascade through food webs, and how trophic structure influences their effects.

### Functional redundancy in carbon utilization potential

Despite the marked changes in taxonomic composition driven by glyphosate, microbial communities did not change their carbon substrate use throughout the experiment, providing evidence for functional redundancy in carbon utilization potential. This was an expected result, as broad-scale ecosystem functions such as respiration and dissolved organic carbon consumption are weakly coupled with species composition (Girvan, Campbell, Killham, Prosser, & Glover, 2005; Langenheder, Lindström, & Tranvik, 2006; Peter et al., 2011), allowing these functions to remain unaffected by fluctuations in microbial community composition (Louca et al., 2018). While less diverse communities (in terms of species richness) may lack functional redundancy, more diverse communities are expected to encode more redundant functions (Konopka, 2009). We can thus surmise that the freshwater bacterioplankton communities studied here were sufficiently diverse to be functionally redundant for carbon utilization in the face of disturbance. The weak and time-independent effect of high concentrations of glyphosate on alpha diversity was insufficient to alter community carbon substrate use. However, our experiment was conducted with communities originating from a pristine lake in a nature reserve, and this result might not be generalized to freshwaters historically impacted by other forms of anthropogenic stress. For example, land use intensity is negatively correlated with bacterioplankton richness in lakes across Eastern Canada (Kraemer et al., 2020). It remains to be seen whether such impacted lakes are less functionally redundant, and thus possibly more susceptible to impaired ecosystem functioning. Lastly, although bacterioplankton respiration accounts for a large fraction of organic carbon processing in freshwaters (Berggren, Lapierre, & del Giorgio, 2012), the carbon substrate use we measured could also be due in part to fungal activity which could be compensating or masking changes in bacterioplankton activity. There was no macroscopically observable fungal growth in the plates, yet microscopic fungi likely contributed a fraction of the inoculum used to initiate the plates.

### The phylogenetic depth of glyphosate resistance: Methodological considerations

The inference of bacterioplankton ASVs in this study allowed a relatively fine-scale taxonomic resolution of community changes in response to a pulse perturbation of glyphosate. Notably, the recovery of bacterioplankton composition was detectable at broader taxonomic units (e.g. phylum in particular) but not at the ASV level. This implies that the taxonomic resolution of traits under selection during recovery from a glyphosate pulse is relatively coarse (Martiny, Jones, Lennon, & Martiny, 2015). This result could also be explained if ASVs are too fine-scale as a measure of diversity, and mostly reflect sequencing or base calling errors rather than true biological diversity. We deem this unlikely, first because the ASV inference algorithm includes a model-based approach to correct for amplicon sequencing errors (Callahan et al., 2016), and second because ASV detection methods are usually more accurate than OTU-clustering methods based on sequence similarity thresholds of usually 97% (Caruso, Song, Asquith, & Karstens, 2019). For example, we only found 7 to 10 false-positive ASVs (Methods), but dozens to hundreds of false positive are detected by even state-of-the-art (distribution-based) sequence clustering-based methods to identify operational taxonomic units, when applied to the same or similar mock communities as used here (Tromas et al., 2017).

Although we cannot exclude the impact of possible false ASVs on our results, we expect them to be relatively minimal and evenly distributed across all timepoints (Callahan, McMurdie, & Holmes, 2017). In other words, there is no reason to believe that sequencing errors should be non-randomly distributed over time or across experimental treatments. Moreover, PRC analyses show a steady decline from the phylum level to the genus level in both the response to, and recovery from, high concentrations of glyphosate. Therefore, even without considering the ASV level, there is still a discernible pattern of greater community resilience at broader taxonomic scales. This suggests that the traits (and underlying genes) required for survival or growth in the presence of glyphosate are relatively deeply conserved. Higher-resolution genomic or metagenomic analyses could be used to confirm this result, and pinpoint the genes involved in resistance.

### Ecosystem resistance, resilience and stability

Our study provides evidence of ecosystem stability in terms of carbon substrate use maintained by microbial communities when faced by a perturbation by two of the most commonly used pesticides in the world, separately or in combination. We also showed resistance to a wide gradient of imidacloprid contamination, and resilience to high doses of glyphosate in bacterioplankton communities that have no known history of contact with the herbicide. Finally, whether a stressed community is considered resilient depends on the phylogenetic depth of the traits required to deal with the stress (Martiny et al., 2015). Our results provide an example of how resilience to stressors can be a feature of deeper phylogenetic groups, but not finer-scale groupings (ASVs), which could be involved in adaptation to other stressors or niches.

## Acknowledgments

We are grateful to D. Maneli, C. Normandin, A. Arkilanian and T. Jagadeesh for their assistance in the field, to J. Marleau, C. Girard, O.M. Pérez-Carrascal and N. Tromas for their assistance during laboratory analyses, to K. Velghe for nutrient analyses and to M.A.P. Castro for developing the LC–MS method for pesticides quantification. We thank the three anonymous reviewers for comments on a previous version of this manuscript.

## Funding information

This study was supported by a Canada Research Chair and NSERC Discovery Grant to B.J.S. N.B.C. was funded by FRQNT and NSERC-CREATE/GRIL fellowships. V.F. was supported by an NSERC postdoctoral fellowship. M-P.H. was funded by NSERC and NSERC-CREATE/GRIL. C.C.Y.X. was funded by a Vanier Canada Graduate Scholarship. R.D.H.B. was supported by a Canada Research Chair. LEAP was built and operated with funds from a CFI Leaders Opportunity Fund, NSERC Discovery Grant and the Liber Ero Chair to A.G.

## Data accessibility

Sequence data corresponding to raw 16S rRNA reads and metadata, as well as carbon substrate utilization dataset based on Biolog EcoPlates assessment, are available on https://figshare.com/projects/MEC-LEAP/78297. Sequences of 16S rRNA reads are also available at NCBI SRA (BioProject ID PRJNA664121).

## Author contributions

N.B.C., V.F., M.-P.H., R.D.H.B., B.E.B., G.B., G.F.F., B.J.S. and A.G. designed the study. N.B.C., V.F. and M.-P.H. collected the data. N.B.C., C.C.Y.X. and V. Y. contributed to the development of laboratory methods. N.B.C. and V.F. analysed data. N.B.C. made the figures and drafted the manuscript. N.B.C. and B.J.S. wrote the first manuscript draft and all authors contributed significantly to data interpretation and commented on manuscript drafts.

## Supplementary figures captions

**Fig. S1** Experimental gradient established for (A) glyphosate and (B) imidacloprid concentrations between two application pulses (at days 6 and 34) and (C) the correlation between target and measured concentrations at each pulse. The top row of figure C shows results for glyphosate, and the bottom two rows for imidacloprid, after pulse 1 (left column) and pulse 2 (right column) respectively.

**Fig. S2** PRC plots show no effect of experimental treatments on community metabolic profiles when considering (A) the 31 different compounds individually (F=32.6 p=0.69) or (B) grouped according to functional guilds (F=79.2 p=0.86). The PRC axis shown in A explains 13.4% of total variance and in B 43.1%.

**Fig. S3** PCoA ordinations based on (A, B) weighted UniFrac distance or (B, D) Jensen-Shannon divergence exploring different normalization approaches: (A, C) calculation of reads relative abundance and (B, D) rarefying to a threshold of 10,000 reads per sample. Each sampling day is plotted separately to facilitate visualization of treatment effects on community composition.

**Fig. S4** PRCs show how high glyphosate treatments affected community composition at (A) phylum and (B) ASV levels. Low- and high-nutrient treatments show the same pattern, and, for this reason, they were grouped in Fig. 4, to facilitate data visualization. The finest level of taxonomic assignment based on FreshTrain and GreenGenes database is shown following ASV names in panel B. Only taxa with weights higher than 0.2 are shown in A and higher than 0.095 are shown in B.

**Fig. S5** PRCs show how high glyphosate treatments (15 mg/L) affected community composition at different taxonomic levels: (A) class, (B) order, (C) family/lineage, (D) genus/clade. Taxonomic assignment based on FreshTrain and GreenGenes databases. Low and high nutrient treatments were grouped as they follow the same pattern. Only taxa with weights higher than 0.2 are shown.

**Fig. S6** Summed effects of GAMMs on abundance of three genera most positively affected by the glyphosate treatments: (A) Agrobacterium, (B) Flavobacterium and (C) Azospirillum. Shades indicate a confidence interval of 95%. Abundance of each genus is the estimated absolute abundance of all ASVs assigned to Agrobacterium, Flavobacterium or Azospirillum after normalization by rarefying each sample to 10,000 reads without replacement.

## Supplementary tables captions

**Table S1** Regulatory acceptable concentrations (RACs) of glyphosate and imidacloprid residues in freshwater according to regulatory agencies in Canada (CCME, Canadian Council of Ministers of the Environment), Europe (EFSA, European Food Safety Agency) and in the USA (EPA, Environmental Protection Agency). Chronic (long-term) and acute (short-term) exposure RACs are specified when available.

**Table S2** Carbon substrates present in Biolog EcoPlates and their respective grouping (guild)

**Table S3** Barcode sequences of the reverse primer used in step 2 PCR, and total read counts obtained after sample demultiplexing. The number of non-chimeric reads obtained after filtering, denoising, merging paired ends and removing chimeras with DADA2, is also shown.

**Table S4** Summarized results of the generalized additive mixed models (GAMMs) for bacterial density and number of carbon substrate used as a response variables. A Gaussian residual distribution was used for both models. For each response variable we report the sample size (n), adjusted R2, the predictors used in the model, the parameter estimate and respective standard error (SE) of parametric effects or the effective degrees of freedom (EDF) of smooth terms, the corresponding test statistics (*t* value for parametric and F for smooth terms) and the *p*-value. Smooths terms are described as mgcv syntax: ‘s()’ functions are thin plate regression splines and ‘ti()’ tensor product interactions, pond represents the random variable of the mixed model and ‘bs=‘fs’’ specifies the underline base function as a random smooth. Following a Bonferroni multiple testing correction for 9 tests, we only considered significant variables with unadjusted *p*-value <0.005 (shown in bold).

**Table S5** Summarized results of the generalized additive mixed models (GAMMs) for alpha diversity: observed ASV and exponential Shannon. Gaussian residual distributions were used in all models. For each response variable we report the sample size (n), adjusted R2, the predictors and factors used in the model, the parameter estimate and respective standard error (SE) of parametric effects or the effective degrees of freedom (EDF) of smooth terms, the corresponding test statistics (*t* value for parametric and F for smooth terms) and the *p*-value. Smooths terms are described as mgcv syntax: ‘s()’ functions are thin plate regression splines and ‘ti()’ tensor product interactions, pond represents the random variable of the mixed model and ‘bs=‘fs’’ specifies the underline base function as a random smooth. Following a Bonferroni multiple testing correction for 16 tests, we only considered significant variables with *p*-value <0.003, shown in bold.

**Table S6** PERMANOVA for different explanatory variables (and their interaction) in models with the weighted UniFrac distances among communities as the response. The same model was tested at five different time points and an asterisk indicates p-values that are significant after a Bonferroni correction for 7 hypothesis tests (i.e. *p*-values <0.007 are considered significant, shown in bold). A PERMDISP was performed to confirm homogeneity of groups dispersions and significant p-values (<0.05) point out to predictors whose significance in the PERMANOVA output should be carefully analysed as they may be a result of within-group variation rather than among-group variation. df=degrees of freedom

**Table S7** PERMANOVA for different explanatory variables (and their interaction) in models with the Jensen-Shannon divergence among communities as the response. The same model was tested at five different time points and an asterisk indicates p-values that are significant after a Bonferroni correction for 7 hypothesis tests (i.e. only *p*-values <0.007 are considered significant, shown in bold). A PERMDISP was performed to confirm homogeneity of groups dispersions and significant *p*-values (<0.05) point out to predictors whose significance in the PERMANOVA output should be carefully analysed as they may be a result of within-group variation rather than among-group variation. df=degrees of freedom

**Table S8** PERMANOVA for glyphosate as the explanatory variable in models with weighted UniFrac distance or Jensen-Shannon divergence among communities as the response variable after data transformation by ASV relative abundance calculation (unrarefied) or by rarefying samples to 10,000 reads. The same model was tested at five different time points and an asterisk indicates p-values that are significant after a conservative Bonferroni correction for 7 hypothesis tests (i.e. only *p*-value<0.007 are considered significant). A PERMDISP was performed to confirm homogeneity of groups dispersions and significant *p*-values (<0.05) point out to predictors whose significance in the PERMANOVA output should be carefully analysed as they may be a result of within- group variation rather than among-group variation.

**Table S9** Percent of variance explained by the two first PRC axes, and by time or treatment when nutrient treatments are grouped as replicates (see Fig. 6 and Fig. S5). F statistic and *p*-value of permutation test for first constrained eigenvalue is also shown, and an asterisk denote significant *p*-values.

**Table S10** All bacterioplankton taxa weights for the PRC model at the phylum level, ranked from largest (positive effects of glyphosate treatment) to smallest (negative effects of glyphosate treatment).

**Table S11** Relative abundance of the main affected phyla by treatment. Percentage was calculated after normalization with DESeq2 or by rarefying samples to 10,000 reads each and the respective standard error is indicated in parenthesis.

**Table S12** ASVs with the highest PRC taxa weights, and their respective weight in the first RDA axis, the ratio between this value and the maximum taxa weight of the PRC model, and their taxonomy assignment from TaxAss using FreshTrain and GreenGenes databases.

**Table S13** Summarized results of the generalized additive mixed models (GAMMs) for abundance of the three genera most positively impacted by the experimental treatments. Gaussian residual distributions were used in all models. For each response variable we report the sample size (n), adjusted R2, the predictors and factors used in the model, the parameter estimate and respective standard error (SE) of parametric effects or the effective degrees of freedom (EDF) of smooth terms, the corresponding test statistics (*t* value for parametric and F for smooth terms) and the *p*-value. Smooths terms are described as mgcv syntax: ‘s()’ functions are thin plate regression splines and ‘t()’ tensor product interactions, pond represents the random variable of the mixed model and ‘bs=‘fs’’ specifies the underline base function as a random smooth. Following a Bonferroni multiple testing correction for 16 tests, we only considered significant variables with *p*-value <0.003, shown in bold.

## Notes

### Competing Interest Statement

The authors have declared no competing interest.

### Summary of Updates

Significant new data reanalysis, but major conclusions remain unchanged.

