## Supplementary Methods, Figures, and Tables for "Resistance, resilience, and functional redundancy of freshwater bacterioplankton communities facing a gradient of agricultural stressors in a mesocosm experiment"

Table of Contents:

| Supplementary methods | Page 2 |
| --- | --- |
| Table S1 | Page 3 |
| Table S2 | Page 4 |
| Table S3 | Page 5 |
| Table S4 | Page 8 |
| Table S5 | Page 9 |
| Table S6 | Page 11 |
| Table S7 | Page 13 |
| Table S8 | Page 15 |
| Table S9 | Page 16 |
| Table S10 | Page 17 |
| Table S11 | Page 18 |
| Table S12 | Page 19 |
| Table S13 | Page 20 |
| Supplementary figures captions | Page 23 |
| Fig S1 | Page 24 |
| Fig S2 | Page 25 |
| Fig S3 | Page 26 |
| Fig S4 | Page 27 |
| Fig S5 | Page 28 |
| Fig S6 | Page 29 |

**Supplementary methods**

**Illumina library preparation and sequencing**

We performed a two-step PCR amplification of the V4 region of the 16S rRNA gene to first amplify the region (step 1) and then attach barcodes and Illumina adapters to the amplicon (step 2) following protocol described in Preheim et al. (2013). All PCR reactions were performed in Mastercycler nexus thermocyclers (Eppendorf Corporate, Mississauga, Canada).

Step 1 PCR was performed with 0.5 unit of Phusion DNA polymerase and 1X Phusion High Fidelity Buffer (ThermoFisher, Waltham, MA, USA), 200 µl of dNTPs (ThermoFisher, Waltham, MA, USA), 0.36 µM of each primer (U515_F and E786_R) and 20 ng of extracted environmental DNA in reactions of 25 µl. PCR conditions were: initial denaturation for 30 seconds at 98 ˚C, followed by 22 cycles of denaturation at 98 ˚C for 20 seconds, annealing at 54 ˚C for 35 seconds and extension at 72 ˚C for 30 seconds, the cycles were followed by final elongation at 72 ˚C for 60 seconds. Four reactions of 25 µl were performed per sample, pooled and cleaned with the Zymo research DNA purification kit (Zymo Research, Irvine, USA) according to manufacturer’s protocol.

Step 2 PCR was performed similarly to step 1, but with 4 µl of the purified step 1 PCR product and 0.36 µM of PE-III-PCR-F and 0.36 µM of barcoded reverse primers PE-III-PCR-XXXX (see exact barcode sequence of each sample in Table S3). PCR conditions were: initial denaturation for 30 seconds at 98˚C, followed by seven cycles of 98˚C for 30 seconds, 83˚C for 30 seconds and 72˚C for 30 seconds. Reactions were performed in duplicates, then pooled and purified using Agencourt AMPure XP beads (Beckman Coulter Life Sciences, Indianopolis, IN, USA) following manufacturer’s instructions. Fragment size was confirmed via electrophoresis in agarose gels and quantified on a NanoDrop microvolume spectrophotometer (ThermoFisher, Waltham, MA, USA).

Libraries were then pooled at equimolar ratio, denatured and sequenced, using the MiSeq reagent Kit V2 with 500 cycles (Illumina, San Diego, CA, USA) yielding two 250 bp paired-end reads. The 96 samples were split between two different sequencing runs, each of them including PCR negative controls, extraction blanks, and a mock community DNA sample containing identified 16S rRNA clone libraries. PCR negative controls consisted of ultrapure DNase/RNase-free distilled water (ThermoFisher, Waltham, MA, USA), and DNA extraction blanks from clean filters. In one of the runs we used a custom mock community composed of 16S rRNA clone libraries from freshwater lake samples (Preheim et al. 2013) and in the other we used the American Type Culture Collection MSA-1002 mock community (ATCC, Manassas, VA, USA).

**Table S1** Regulatory acceptable concentrations (RACs) of glyphosate and imidacloprid residues in freshwater according to regulatory agencies in Canada (CCME, Canadian Council of Ministers of the Environment), Europe (EFSA, European Food Safety Agency) and in the USA (EPA, Environmental Protection Agency). Chronic (long-term) and acute (short-term) exposure RACs are specified when available.

| Regulatory agency | Glyphosate | Imidacloprid |
| --- | --- | --- |
| CCME (Canada) | 800 µg/L chronic  27,000 µg/L acute | 0.23 µg/L interim |
| EFSA (Europe) | Risk to aquatic organisms considered low  Selected data from toxicological studies:  - 12,500 µg/L chronic toxicity to *Daphnia magna*  - 40,000 µg/L acute toxicity to *D. magna*  - 8,500 µg/L acute toxicity to *Aphanizomenon flosaquae* | Tier 1^†^  - 0.209 µg/L chronic  - 0.341 µg/L acute  Tier-2B^†^  - 0.009 µg/L chronic  - 0.098 µg/L acute |
| EPA (USA) ^‡^ | 26,600 µg/L chronic  49,900 µg/L acute | 0.01 µg/L chronic  0.385 µg/L acute |

^†^ Tier 1 is indicated as not appropriate for risk assessment and Tier-2B to be used provisional risk for assessment

^‡^ EPA's Office of Pesticide Programs (OPP) for aquatic invertebrates

**Table S2** Carbon substrates present in Biolog EcoPlates and their respective grouping (guild)

| Substrate | Group (guild) |
| --- | --- |
| Phenylethylamine | Amines/amides |
| Putrescine | Amines/amides |
| L-arginine | Amino acids |
| L-asparagine | Amino acids |
| L-phenylalanine | Amino acids |
| L-serine | Amino acids |
| L-threonine | Amino acids |
| Glycyl-L-glutamic acid | Amino acids |
| Pyruvic acid methyl ester | Carbohydrates |
| D-cellobiose | Carbohydrates |
| Alpha-D-lactose | Carbohydrates |
| Beta-methyl-D-glucoside | Carbohydrates |
| D-xylose | Carbohydrates |
| i-erythritol | Carbohydrates |
| D-mannitol | Carbohydrates |
| N-acetyl-D-glucosamine | Carbohydrates |
| Glucose-1-phosphate | Carbohydrates |
| D,L-alpha-glycerol phosphate | Carbohydrates |
| D-glucosaminic acid | Carboxylic acetic acids |
| D-galactonic acid-gamma-lactone | Carboxylic acetic acids |
| D-galacturonic acid | Carboxylic acetic acids |
| 2-Hydroxy benzoic acid | Carboxylic acetic acids |
| 4-Hydroxy benzoic acid | Carboxylic acetic acids |
| Gamma-hydroxybutyric acid | Carboxylic acetic acids |
| Itaconic acid | Carboxylic acetic acids |
| Alpha-ketobutyric acid | Carboxylic acetic acids |
| D-malic acid | Carboxylic acetic acids |
| Tween 40 | Polymers |
| Tween 80 | Polymers |
| Alpha-cyclodextrin | Polymers |
| Glycogen | Polymers |

**Table S3** Barcode sequences of the reverse primer used in step 2 PCR, and total read counts obtained after sample demultiplexing. The number of non-chimeric reads obtained after filtering, denoising, merging paired ends and removing chimeras with DADA2, is also shown.

| Sample | Barcode sequence | Total reads | Non-chimeric reads |
| --- | --- | --- | --- |
| C1.17.aug | AACCCGTT | 55762 | 50337 |
| C1.23.aug | CAGCGGCA | 17714 | 15227 |
| C1.28.sep | GTTCGCAG | 14880 | 13926 |
| C1.31.aug | CCGACAAA | 51823 | 44758 |
| C1.15.sep | TAAGGGAG | 12294 | 11295 |
| C1.20.sep | TTGTGGCG | 23611 | 22246 |
| C4.17.aug | GACATCAT | 56234 | 50201 |
| C4.23.aug | GAGTTTGA | 7958 | 7118 |
| C4.31.aug | AACAGTAT | 15241 | 11601 |
| C4.15.sep | ATCGCACC | 39833 | 36356 |
| C4.20.sep | CTAGAATC | 12371 | 7857 |
| C4.28.sep | CCTTTGAT | 23931 | 21410 |
| C8.17.aug | ATAGGTGG | 18402 | 16033 |
| C8.23.aug | CAACTTCA | 29342 | 27344 |
| C8.31.aug | GTAGTCGA | 21294 | 19270 |
| C8.15.sep | TCCCGATG | 19440 | 18054 |
| C8.20.sep | GGGCGAAA | 13574 | 11480 |
| C8.28.sep | GGTGTACC | 16869 | 14077 |
| D1.17.aug | CGTCCCAC | 37000 | 33163 |
| D1.23.aug | CTGTTAGT | 36130 | 33232 |
| D1.31.aug | CACTCACT | 32340 | 29168 |
| D1.15.sep | ACCTCCCA | 43478 | 40627 |
| D1.20.sep | AATACAGG | 66220 | 60120 |
| D1.28.sep | AGTCACCC | 9413 | 8460 |
| D4.17.aug | TACGATAC | 24498 | 21285 |
| D4.23.aug | AGGCTTCA | 10161 | 9333 |
| D4.31.aug | GTGCTGAT | 39070 | 34550 |
| D4.15.sep | ACCATACT | 38941 | 36695 |
| D4.20.sep | AAATTGGA | 44426 | 40952 |
| D4.28.sep | TTCCAGAT | 12791 | 11531 |
| D8.17.aug | GCCTGTTC | 35891 | 32901 |
| D8.23.aug | TCGGCTCG | 85995 | 80978 |
| D8.31.aug | AGAGAGGC | 30980 | 24001 |
| D8.15.sep | GCAATGGA | 29363 | 26644 |
| D8.20.sep | TGACTTAG | 34660 | 31171 |
| D8.28.sep | GGAGGCTG | 21580 | 16647 |
| E1.17.aug | CCGCACCG | 27000 | 24008 |
| E1.23.aug | ATGCCAGC | 15379 | 13891 |
| E1.31.aug | TCGAACAC | 68782 | 63125 |
| E1.15.sep | CGACATTC | 26909 | 20436 |
| E1.20.sep | CATCGCTA | 41282 | 34865 |
| E1.28.sep | ACTAAGAT | 18204 | 15298 |
| E4.17.aug | TCAAAGCT | 70496 | 61775 |
| E4.23.aug | CAGCGGCA | 49463 | 44749 |
| E4.31.aug | CCGACAAA | 55130 | 49059 |
| E4.15.sep | TAAGGGAG | 50914 | 46906 |
| E4.20.sep | TTGTGGCG | 49999 | 44315 |
| E4.28.sep | GTTCGCAG | 47427 | 42091 |
| E8.17.aug | TGGGACCT | 61084 | 52159 |
| E8.23.aug | GAGTTTGA | 28586 | 25272 |
| E8.31.aug | AACAGTAT | 54729 | 48638 |
| E8.15.sep | ATCGCACC | 45603 | 39995 |
| E8.20.sep | CTAGAATC | 36482 | 32974 |
| E8.28.sep | CCTTTGAT | 65278 | 59393 |
| H1.17.aug | TGTTTCCC | 12056 | 10311 |
| H1.23.aug | GGTAATGA | 15470 | 13598 |
| H1.31.aug | GTACGTTG | 48086 | 44387 |
| H1.15.sep | ACGGGCTG | 11257 | 9873 |
| H1.20.sep | ATGAAGTA | 12745 | 11221 |
| H1.28.sep | ACACCTCG | 10727 | 9949 |
| H4.17.aug | ATAGGTGG | 38081 | 35342 |
| H4.23.aug | CAACTTCA | 53637 | 49787 |
| H4.31.aug | GTAGTCGA | 40116 | 36626 |
| H4.15.sep | TCCCGATG | 65142 | 60387 |
| H4.20.sep | GGGCGAAA | 29985 | 27212 |
| H4.28.sep | GGTGTACC | 38098 | 35100 |
| H8.17.aug | CGTCCCAC | 71142 | 62412 |
| H8.23.aug | CTGTTAGT | 64022 | 57820 |
| H8.31.aug | CACTCACT | 76152 | 66286 |
| H8.15.sep | ACCTCCCA | 72837 | 66412 |
| H8.20.sep | GAGCACAG | 84816 | 77740 |
| H8.28.sep | CGAATATT | 65034 | 58318 |
| J4.17.aug | TACGATAC | 75103 | 67734 |
| J4.23.aug | AGGCTTCA | 54008 | 48385 |
| J4.31.aug | GTGCTGAT | 54083 | 50492 |
| J4.15.sep | ACCATACT | 62067 | 57735 |
| J4.20.sep | AAATTGGA | 65309 | 59721 |
| J4.28.sep | TTCCAGAT | 30185 | 27734 |
| J8.17.aug | GCCTGTTC | 72714 | 63613 |
| J8.23.aug | AGCTGACG | 58664 | 50262 |
| J8.31.aug | AGAGAGGC | 55708 | 48142 |
| J8.15.sep | GCAATGGA | 58161 | 52962 |
| J8.20.sep | TGACTTAG | 64244 | 56997 |
| J8.28.sep | GGAGGCTG | 43352 | 37922 |
| K4.17.aug | CCGCACCG | 80953 | 74716 |
| K4.23.aug | ATGCCAGC | 81279 | 74020 |
| K4.31.aug | TCGAACAC | 96033 | 88819 |
| K4.15.sep | CGACATTC | 44684 | 41105 |
| K4.20.sep | CATCGCTA | 48869 | 44067 |
| K4.28.sep | ACTAAGAT | 64643 | 59626 |
| K8.17.aug | TGTTTCCC | 80622 | 73294 |
| K8.23.aug | GGTAATGA | 76790 | 69163 |
| K8.31.aug | GAAACTGG | 30648 | 27342 |
| K8.15.sep | ACGGGCTG | 45460 | 42448 |
| K8.20.sep | ATGAAGTA | 42178 | 37273 |
| K8.28.sep | ACACCTCG | 51882 | 45344 |

**Table S4** – Summarized results of the generalized additive mixed models (GAMMs) for bacterial density and number of carbon substrate used as a response variables. A Gaussian residual distribution was used for both models. For each response variable we report the sample size (n), adjusted R^2^, the predictors used in the model, the parameter estimate and respective standard error (SE) of parametric effects or the effective degrees of freedom (EDF) of smooth terms, the corresponding test statistics (*t* value for parametric and F for smooth terms) and the *p*-value. Smooths terms are described as mgcv syntax: ‘s()’ functions are thin plate regression splines and ‘ti()’ tensor product interactions, pond represents the random variable of the mixed model and ‘bs=‘fs’’ specifies the underline base function as a random smooth. Following a Bonferroni multiple testing correction for 9 tests, we only considered significant variables with unadjusted p-value <0.005 (shown in bold).

| Response variable | n | Adjusted R^2^ | Predictors (significant effects in bold) | Estimate (SE) or EDF | Statistic | *p*-value |
| --- | --- | --- | --- | --- | --- | --- |
| log_10_(Bacterial density) | 288 | 0.734 | **nutrient**  **ti(day)**  **ti(day, glyphosate)**  ti(day, glyphosate, by = nutrient)  ti(day, imidacloprid)  ti(day, imidacloprid, by = nutrient)  ti(day, glyphosate, imidacloprid)  ti(day, glyphosate, imidacloprid, by = nutrient)  **s(day, Pond, bs='fs')** | **0.08 (SE=0.02)**  **4.6**  **4.1**  1.0  2.4  1.0  0.0  3.1  **70.5** | **4.1**  **17.5**  **6.6**  5.7  0.2  0.5  0.0  2.1  **2.2** | **< 0.001**  **< 0.001**  **< 0.001**  0.018  0.916  0.490  0.183  0.260  **< 0.001** |
| Number of carbon substrates used (EcoPlates) | 288 | 0.309 | nutrient  **ti(day)**  ti(day, glyphosate)  ti(day, glyphosate, by = nutrient)  ti(day, imidacloprid)  ti(day, imidacloprid, by = nutrient)  ti(day, glyphosate, imidacloprid)  ti(day, glyphosate, imidacloprid, by = nutrient)  **s(day, Pond, bs='fs')** | 0.88 (SE=0.37)  **4.5**  1.0  3.1  1.0  6.0  0.0  1.6  **33.4** | 2.4  **6.0**  1.0  1.8  1.4  1.3  0.0  1.1  **0.4** | 0.019  **< 0.001**  0.320  0.136  0.242  0.244  0.266  0.449  **< 0.001** |

**Table S5** – Summarized results of the generalized additive mixed models (GAMMs) for alpha diversity: observed ASV and exponential Shannon. Gaussian residual distributions were used in all models. For each response variable we report the sample size (n), adjusted R^2^, the predictors and factors used in the model, the parameter estimate and respective standard error (SE) of parametric effects or the effective degrees of freedom (EDF) of smooth terms, the corresponding test statistics (*t* value for parametric and F for smooth terms) and the *p*-value. Smooths terms are described as mgcv syntax: ‘s()’ functions are thin plate regression splines and ‘ti()’ tensor product interactions, pond represents the random variable of the mixed model and ‘bs=‘fs’’ specifies the underline base function as a random smooth. Following a Bonferroni multiple testing correction for 16 tests, we only considered significant variables with *p*-value <0.003, shown in bold.

| Response variable | n | Adjusted R^2^ | Predictors | Factors^†^ of parametric and smooth terms | Estimate (SE) or EDF | Statistic | *p*-value |
| --- | --- | --- | --- | --- | --- | --- | --- |
| log_10_(Observed ASVs) | 89 | 0.69 | treatment  nutrient  ti(day)  ti(day, by=treatment)  ti(day, by=nutrient)  **s(day, Pond, bs='fs')** | high_both  high_glypho  high_imid  low_both  low_glypho  low_imid  high  low_glypho  low_imid  low_both  high_glypho  high_imid  high_both  high | -0.02 (0.07)  -0.20 (0.07)  0.07 (0.07)  0.13 (0.07)  -0.02 (0.07)  0.14 (0.07)  -0.09 (0.04)  4.7  2.4  1.0  1.0  2.0  1.0  3.6  11.2  **1.4** | -0.27  -2.89  1.06  1.88  -0.26  2.03  2.21  3.18  0.48  0.51  0.21  0.94  0.51  2.49  1.34  **1.17** | 0.786  0.006  0.296  0.065  0.795  0.048  0.032  0.021  0.589  0.509  0.652  0.539  0.478  0.110  0.372  **<0.001*** |
| exp(Shannon index) | 89 | 0.66 | **treatment**  nutrient  ti(day)  ti(day, by=treatment)  ti(day, by=nutrient)  **s(day, Pond, bs='fs')** | high_both  **high_glypho**  high_imid  low_both  low_glypho  low_imid  high  low_glypho  low_imid  low_both  high_glypho  high_imid  high_both  high | -7.3 (4.3)  **-14.9 (4.2)**  -5.1 (4.2)  4.1 (4.3)  1.2 (4.4)  11.4 (4.3)  0.9 (2.5)  4.0  1.0  3.7  2.5  1.7  1.0  3.3  2.2  **10.2** | -1.70  **-3.51**  -1.22  0.95  0.28  2.64  0.37  1.62  1.95  3.05  0.46  0.77  0.00  4.36  2.60  **1.00** | 0.096  **0.001***  0.228  0.346  0.780  0.011  0.715  0.262  0.169  0.018  0.706  0.526  0.993  0.012  0.116  **0.001*** |

^†^ When factor is absent it means the respective predictor variable is continuous (“day”) or a random effect (“pond”).

**Table S6** – PERMANOVA for different explanatory variables (and their interaction) in models with the weighted UniFrac distances among communities as the response. The same model was tested at five different time points and an asterisk indicates *p*-values that are significant after a Bonferroni correction for 7 hypothesis tests (i.e*.* *p*-values <0.007 are considered significant, shown in bold). A PERMDISP was performed to confirm homogeneity of groups dispersions and significant *p*-values (<0.05) point out to predictors whose significance in the PERMANOVA output should be carefully analysed as they may be a result of within-group variation rather than among-group variation. df=degrees of freedom

|  |  |  | PERMANOVA  (999 permutations) | | | PERMDISP  (999 permutations) | |
| --- | --- | --- | --- | --- | --- | --- | --- |
|  | Predictor | df | F | R^2^ | *p*-value | F | *p*-value |
| Day 1 | Nutrients | 1 | 1.02 | 0.09 | 0.445 | 0.04 | 0.855 |
|  | Glyphosate | 2 | 0.59 | 0.10 | 0.925 | 0.94 | 0.412 |
|  | Imidacloprid | 2 | 0.76 | 0.13 | 0.775 | 1.86 | 0.182 |
|  | Nutrients and glyphosate | 2 | 0.63 | 0.11 | 0.894 |  |  |
|  | Nutrients and imidacloprid | 2 | 0.49 | 0.09 | 0.979 |  |  |
|  | Glyphosate and imidacloprid | 2 | 1.15 | 0.20 | 0.362 |  |  |
|  | Nutrients, glyphosate and imidacloprid | 2 | 0.59 | 0.10 | 0.937 |  |  |
| Day 7 | Nutrients | 1 | 2.65 | 0.07 | 0.131 | 0.02 | 0.873 |
|  | **Glyphosate** | **2** | **10.40** | **0.58** | ***0.005**** | 0.29 | 0.754 |
|  | Imidacloprid | 2 | 1.13 | 0.06 | 0.429 | 3.21 | 0.065 |
|  | Nutrients and glyphosate | 2 | 1.24 | 0.07 | 0.378 |  |  |
|  | Nutrients and imidacloprid | 2 | 1.06 | 0.06 | 0.461 |  |  |
|  | Glyphosate and imidacloprid | 2 | 0.96 | 0.05 | 0.472 |  |  |
|  | Nutrients, glyphosate and imidacloprid | 2 | 0.80 | 0.04 | 0.572 |  |  |
| Day 15 | Nutrients | 1 | 4.75 | 0.12 | 0.027 | 0.11 | 0.701 |
|  | **Glyphosate** | **2** | **10.06** | **0.53** | ***0.001**** | 0.67 | 0.523 |
|  | Imidacloprid | 2 | 0.91 | 0.05 | 0.537 | 0.93 | 0.930 |
|  | Nutrients and glyphosate | 2 | 1.95 | 0.10 | 0.169 |  |  |
|  | Nutrients and imidacloprid | 2 | 0.92 | 0.05 | 0.532 |  |  |
|  | Glyphosate and imidacloprid | 2 | 1.12 | 0.06 | 0.417 |  |  |
|  | Nutrients, glyphosate and imidacloprid | 2 | 0.83 | 0.04 | 0.596 |  |  |
| Day 30 | Nutrients | 1 | 2.44 | 0.12 | 0.055 | 0.04 | 0.848 |
|  | Glyphosate | 2 | 2.92 | 0.29 | 0.007 | 2.06 | 0.169 |
|  | Imidacloprid | 2 | 1.11 | 0.11 | 0.365 | 0.84 | 0.425 |
|  | Nutrients and glyphosate | 2 | 1.11 | 0.11 | 0.383 |  |  |
|  | Nutrients and imidacloprid | 2 | 1.19 | 0.12 | 0.300 |  |  |
|  | Glyphosate and imidacloprid | 2 | 0.89 | 0.09 | 0.590 |  |  |
|  | Nutrients, glyphosate and imidacloprid | 2 | 0.71 | 0.07 | 0.792 |  |  |
| Day 43 | Nutrients | 1 | 3.29 | 0.11 | 0.036 | 0.09 | 0.799 |
|  | **Glyphosate** | **2** | **6.42** | **0.44** | ***0.001**** | 6.05 | 0.015* |
|  | Imidacloprid | 2 | 1.11 | 0.08 | 0.402 | 0.13 | 0.877 |
|  | Nutrients and glyphosate | 2 | 1.55 | 0.10 | 0.217 |  |  |
|  | Nutrients and imidacloprid | 2 | 1.32 | 0.09 | 0.311 |  |  |
|  | Glyphosate and imidacloprid | 2 | 0.89 | 0.06 | 0.543 |  |  |
|  | Nutrients, glyphosate and imidacloprid | 2 | 0.81 | 0.05 | 0.637 |  |  |

**Table S7** – PERMANOVA for different explanatory variables (and their interaction) in models with the Jensen-Shannon divergence among communities as the response. The same model was tested at five different time points and an asterisk indicates *p*-values that are significant after a Bonferroni correction for 7 hypothesis tests (i.e. only *p*-values<0.007 are considered significant, shown in bold). A PERMDISP was performed to confirm homogeneity of groups dispersions and significant *p*-values (<0.05) point out to predictors whose significance in the PERMANOVA output should be carefully analysed as they may be a result of within-group variation rather than among-group variation. df=degrees of freedom

|  |  |  | PERMANOVA  (999 permutations) | | | PERMDISP  (999 permutations) | |
| --- | --- | --- | --- | --- | --- | --- | --- |
|  | Predictor | df | F | R^2^ | *p*-value | F | *p*-value |
| Day 1 | Nutrients | 1 | 1.37 | 0.11 | 0.166 | 0.26 | 0.260 |
|  | Glyphosate | 2 | 0.55 | 0.09 | 0.972 | 0.57 | 0.592 |
|  | Imidacloprid | 2 | 0.90 | 0.15 | 0.614 | 0.78 | 0.500 |
|  | Nutrients and glyphosate | 2 | 0.71 | 0.11 | 0.874 |  |  |
|  | Nutrients and imidacloprid | 2 | 0.52 | 0.08 | 0.990 |  |  |
|  | Glyphosate and imidacloprid | 2 | 1.11 | 0.18 | 0.330 |  |  |
|  | Nutrients, glyphosate and imidacloprid | 2 | 0.69 | 0.11 | 0.883 |  |  |
| Day 7 | Nutrients | 1 | 2.03 | 0.11 | 0.080 | 0.16 | 0.701 |
|  | **Glyphosate** | **2** | **3.28** | **0.34** | ***0.005**** | 1.52 | 0.269 |
|  | Imidacloprid | 2 | 1.21 | 0.13 | 0.278 | 3.54 | 0.057 |
|  | Nutrients and glyphosate | 2 | 0.93 | 0.10 | 0.512 |  |  |
|  | Nutrients and imidacloprid | 2 | 0.72 | 0.07 | 0.776 |  |  |
|  | Glyphosate and imidacloprid | 2 | 0.87 | 0.09 | 0.579 |  |  |
|  | Nutrients, glyphosate and imidacloprid | 2 | 0.63 | 0.07 | 0.858 |  |  |
| Day 15 | Nutrients | 1 | 3.29 | 0.12 | 0.027 | 0.00 | 0.979 |
|  | **Glyphosate** | **2** | **6.37** | **0.45** | ***0.001**** | 2.13 | 0.165 |
|  | Imidacloprid | 2 | 0.91 | 0.06 | 0.526 | 1.27 | 0.305 |
|  | Nutrients and glyphosate | 2 | 1.73 | 0.12 | 0.136 |  |  |
|  | Nutrients and imidacloprid | 2 | 0.80 | 0.06 | 0.621 |  |  |
|  | Glyphosate and imidacloprid | 2 | 1.14 | 0.08 | 0.359 |  |  |
|  | Nutrients, glyphosate and imidacloprid | 2 | 0.68 | 0.05 | 0.759 |  |  |
| Day 30 | Nutrients | 1 | 2.99 | 0.12 | 0.013 | 0.12 | 0.737 |
|  | **Glyphosate** | **2** | **4.21** | **0.34** | ***0.001**** | 8.02 | 0.004* |
|  | Imidacloprid | 2 | 1.48 | 0.12 | 0.125 | 1.02 | 0.378 |
|  | Nutrients and glyphosate | 2 | 1.04 | 0.08 | 0.409 |  |  |
|  | Nutrients and imidacloprid | 2 | 1.07 | 0.09 | 0.385 |  |  |
|  | Glyphosate and imidacloprid | 2 | 1.18 | 0.09 | 0.297 |  |  |
|  | Nutrients, glyphosate and imidacloprid | 2 | 0.98 | 0.08 | 0.479 |  |  |
| Day 43 | Nutrients | 1 | 2.30 | 0.11 | 0.051 | 0.38 | 0.813 |
|  | **Glyphosate** | **2** | **3.85** | **0.37** | ***0.001**** | 10.41 | 0.004* |
|  | Imidacloprid | 2 | 0.91 | 0.09 | 0.549 | 1.14 | 0.345 |
|  | Nutrients and glyphosate | 2 | 0.96 | 0.09 | 0.535 |  |  |
|  | Nutrients and imidacloprid | 2 | 1.01 | 0.10 | 0.446 |  |  |
|  | Glyphosate and imidacloprid | 2 | 0.76 | 0.07 | 0.739 |  |  |
|  | Nutrients, glyphosate and imidacloprid | 2 | 0.78 | 0.07 | 0.699 |  |  |

**Table S8** - PERMANOVA for glyphosate as the explanatory variable in models with weighted UniFrac distance or Jensen-Shannon divergence among communities as the response variable after data transformation by ASV relative abundance calculation (unrarefied) or by rarefying samples to 10,000 reads. The same model was tested at five different time points and an asterisk indicates *p*-values that are significant after a conservative Bonferroni correction for 7 hypothesis tests (i.e. only *p-*value<0.007 are considered significant). A PERMDISP was performed to confirm homogeneity of groups dispersions and significant *p*-values (<0.05) point out to predictors whose significance in the PERMANOVA output should be carefully analysed as they may be a result of within-group variation rather than among-group variation.

|  |  |  | Glyphosate effect (PERMANOVA 999 permutations) | | | Dispersion within glyphosate groups (PERMDISP 999 permutations) | |
| --- | --- | --- | --- | --- | --- | --- | --- |
|  |  | Data transformation | F | R^2^ | *p*-value | F | *p*-value |
| **Weighted UniFrac distance** | Day 1 | Relative abundance | 1.23 | 0.10 | 0.357 | 0.95 | 0.420 |
|  |  | Reads rarefied to 10,000 | 1.24 | 0.10 | 0.358 | 0.71 | 0.522 |
|  | Day 7 | Relative abundance | 3.68 | 0.41 | 0.040 | 0.52 | 0.585 |
|  |  | Reads rarefied to 10,000 | 3.31 | 0.42 | 0.060 | 0.74 | 0.495 |
|  | Day 15 | Relative abundance | **6.06** | **0.33** | ***0.005**** | 0.01 | 0.991 |
|  |  | Reads rarefied to 10,000 | **6.00** | **0.33** | ***0.002**** | 0.00 | 0.996 |
|  | Day 30 | Relative abundance | 2.10 | 0.19 | 0.060 | 0.70 | 0.505 |
|  |  | Reads rarefied to 10,000 | 1.48 | 0.21 | 0.239 | 0.64 | 0.518 |
|  | Day 43 | Relative abundance | 2.83 | 0.36 | 0.031 | **7.14** | ***0.012**** |
|  |  | Reads rarefied to 10,000 | 4.11 | 0.44 | 0.045 | **4.94** | ***0.035**** |
| **Jensen-Shannon divergence** | Day 1 | Relative abundance | 0.65 | 0.08 | 0.831 | 0.60 | 0.522 |
|  |  | Reads rarefied to 10,000 | 0.65 | 0.08 | 0.833 | 0.66 | 0.530 |
|  | Day 7 | Relative abundance | 3.36 | 0.38 | 0.039 | 0.14 | 0.881 |
|  |  | Reads rarefied to 10,000 | 3.10 | 0.40 | 0.059 | 0.43 | 0.682 |
|  | Day 15 | Relative abundance | **6.66** | **0.37** | ***0.002**** | 0.13 | 0.863 |
|  |  | Reads rarefied to 10,000 | **6.59** | **0.37** | ***0.002**** | 0.14 | 0.872 |
|  | Day 30 | Relative abundance | 3.92 | 0.29 | 0.007 | 1.41 | 0.273 |
|  |  | Reads rarefied to 10,000 | 2.87 | 0.30 | 0.056 | 1.23 | 0.298 |
|  | Day 43 | Relative abundance | **3.54** | **0.36** | ***0.005**** | **4.51** | ***0.035**** |
|  |  | Reads rarefied to 10,000 | 3.23 | 0.41 | 0.036 | 1.90 | 0.189 |

**Table S9** – Percent of variance explained by the two first PRC axes, and by time or treatment when nutrient treatments are grouped as replicates (see Fig. 6 and Fig. S5). F statistic and p-value of permutation test for first constrained eigenvalue is also shown, and an asterisk denote significant *p*-values.

| **Taxonomic level** | **PRC1 (%)** | **PRC2 (%)** | **Time (%)** | **Time and treatment (%)** | **Permutation test (999 permutations)** | |
| --- | --- | --- | --- | --- | --- | --- |
|  |  |  |  |  | F | *p*-value |
| Phylum | 47.7 | 15.3 | 15.1 | 46.5 | 31.22 | 0.001* |
| Class | 46.4 | 12.6 | 13.8 | 49.8 | 34.28 | 0.001* |
| Order | 38.6 | 14.5 | 15.3 | 47.2 | 26.19 | 0.001* |
| Family | 33.5 | 11.9 | 16.3 | 45.2 | 21.30 | 0.001* |
| Genus | 34.9 | 10.6 | 14.4 | 44.7 | 20.6 | 0.001* |
| ASV | 22.1 | 8.0 | 11.1 | 41.9 | 10.61 | 0.001* |

**Table S10** – All bacterioplankton taxa weights for the PRC model at the phylum level, ranked from largest (positive effects of glyphosate treatment) to smallest (negative effects of glyphosate treatment).

| **Phylum** | **Weight in PRC1** |
| --- | --- |
| Proteobacteria | 0.37 |
| Chlamydiae | 0.07 |
| Acidobacteria | 0.04 |
| GN02 | 0.02 |
| Firmicutes | 0.01 |
| BRC1 | 0.01 |
| SR1 | 0.004 |
| Fibrobacteres | 2.33E-27 |
| Lentisphaerae | 2.17E-36 |
| Nitrospirae | 4.23E-39 |
| WS3 | 0.00 |
| OP3 | -3.23E-44 |
| Fusobacteria | -1.14E-30 |
| Tenericutes | -0.002 |
| TM6 | -0.03 |
| [Thermi] | -0.03 |
| Gemmatimonadetes | -0.04 |
| TM7 | -0.05 |
| Spirochaetes | -0.05 |
| Chlorobi | -0.06 |
| Armatimonadetes | -0.08 |
| Chloroflexi | -0.08 |
| OD1 | -0.09 |
| Verrucomicrobia | -0.12 |
| Planctomycetes | -0.14 |
| Cyanobacteria | -0.24 |
| Actinobacteria | -0.26 |
| Bacteroidetes | -0.43 |

**Table S11** Relative abundance of the main affected phyla by treatment. Percentage was calculated after normalization with DESeq2 or by rarefying samples to 10,000 reads each and the respective standard error is indicated in parenthesis.

| Normalization | Treatment | Proteobacteria | Bacteroidetes | Actinobacteria | Cyanobacteria |
| --- | --- | --- | --- | --- | --- |
| DESeq2 | Glyphosate 15 mg/L and imidacloprid 60 ug/L | 62.1% (3.2) | 11.7% (2.1) | 2.6% (0.8) | 3.3% (0.8) |
|  | Glyphosate 15 mg/L | 68.2% (3.7) | 10.3% (1.8) | 3.9% (1.5) | 1.6% (0.6) |
|  | Imidacloprid 60 ug/L | 48.4% (1.4) | 20.7% (0.9) | 7.0% (1.0) | 3.6% (0.6) |
|  | Glyphosate 0.3 mg/L and imidacloprid 1 ug/L | 46.4% (0.9) | 20.1% (0.9) | 5.9% (0.8) | 4.0% (0.6) |
|  | Glyphosate 0.3 mg/L | 46.9% (1.4) | 20.6% (1.4) | 6.5% (1.2) | 4.6% (1.7) |
|  | Imidacloprid 1 ug/L | 48.9% (1.3) | 20.1% (1.4) | 6.4% (1.1) | 3.4% (0.5) |
|  | Control | 49.6% (1.2) | 22.4% (1.1) | 5.2% (0.8) | 3.4% (0.6) |
| Rarefied to 10k reads | Glyphosate 15 mg/L and imidacloprid 60 ug/L | 69.5% (5.7) | 12.6% (3.9) | 3.2% (1.6) | 1.1% (0.4) |
|  | Glyphosate 15 mg/L | 66.7% (7.7) | 8.5% (2.5) | 9.5% (5.5) | 0.7% (0.5) |
|  | Imidacloprid 60 ug/L | 52.8% (4.0) | 17.8% (3.9) | 19.9% (4.4) | 1.2% (0.4) |
|  | Glyphosate 0.3 mg/L and imidacloprid 1 ug/L | 38.6% (2.8) | 20.3% (2.6) | 15.8% (4.2) | 2.9% (1.6) |
|  | Glyphosate 0.3 mg/L | 39.8% (3.4) | 16.5% (4.4) | 14.4% (5.7) | 8.1% (4.9) |
|  | Imidacloprid 1 ug/L | 44.3% (1.7) | 24.9% (2.7) | 15.3% (4.2) | 1.9% (0.5) |
|  | Control | 44.4% (4.1) | 20.3% (2.5) | 12.7% (3.9) | 2.2% (0.8) |

**Table S12** – ASVs with the highest PRC taxa weights, and their respective weight in the first RDA axis, the ratio between this value and the maximum taxa weight of the PRC model, and their taxonomy assignment from TaxAss using FreshTrain and GreenGenes databases.

| ASV | Weight in RDA1 | (Taxa weight) / (max. weight) | Phylum | Class | Order | Family/Lineage | Genus/Clade | Species/Tribe |
| --- | --- | --- | --- | --- | --- | --- | --- | --- |
| sp2262 | 0.14 | 1.00 | Proteobacteria | Alphaproteobacteria | unclassified | unclassified | unclassified | unclassified |
| sp188 | 0.13 | 0.92 | Proteobacteria | Betaproteobacteria | Burkholderiales | betI | betI-A | Lhab-A4 |
| sp1895 | 0.12 | 0.88 | Proteobacteria | Alphaproteobacteria | Rhodospirillales | Rhodospirillaceae | *Azospirillum* | *A. massiliensis* |
| sp2118 | 0.12 | 0.84 | Proteobacteria | Alphaproteobacteria | Rhodobacterales | alfVI | unclassified | unclassified |
| sp2155 | 0.11 | 0.81 | Proteobacteria | Alphaproteobacteria | Rhizobiales | Rhizobiaceae | *Agrobacterium* | unclassified |
| sp307 | 0.11 | 0.81 | Proteobacteria | Gammaproteobacteria | Xanthomonadales | Sinobacteraceae | *Nevskia* | *N. ramosa* |
| sp284 | 0.10 | 0.75 | Proteobacteria | Betaproteobacteria | Burkholderiales | betI | betI-A | unclassified |
| sp130 | 0.10 | 0.74 | Proteobacteria | Betaproteobacteria | Methylophilales | betIV | betIV-A | unclassified |
| sp283 | -0.11 | -0.80 | Proteobacteria | Betaproteobacteria | Burkholderiales | betI | betI-A | unclassified |
| sp2111 | -0.11 | -0.82 | Proteobacteria | Alphaproteobacteria | Rhodobacterales | Rhodobacteraceae | *Rhodobacter* | unclassified |

**Table S13** Summarized results of the generalized additive mixed models (GAMMs) for abundance of the three genera most positively impacted by the experimental treatments. Gaussian residual distributions were used in all models. For each response variable we report the sample size (n), adjusted R^2^, the predictors and factors used in the model, the parameter estimate and respective standard error (SE) of parametric effects or the effective degrees of freedom (EDF) of smooth terms, the corresponding test statistics (*t* value for parametric and F for smooth terms) and the *p*-value. Smooths terms are described as mgcv syntax: ‘s()’ functions are thin plate regression splines and ‘t()’ tensor product interactions, pond represents the random variable of the mixed model and ‘bs=‘fs’’ specifies the underline base function as a random smooth. Following a Bonferroni multiple testing correction for 16 tests, we only considered significant variables with *p*-value <0.003, shown in bold.

| Response variable | n | Adjusted R^2^ | Predictors | Factors^†^ of parametric and smooth terms | Estimate (SE) or EDF | Statistic | p-value |
| --- | --- | --- | --- | --- | --- | --- | --- |
| log_10_(*Agrobacterium*) | 89 | 0.80 | **treatment**  nutrient  ti(day)  **ti(day, by=treatment)**  ti(day, by=nutrient)  s(day, Pond, bs='fs') | **high_both**  **high_glypho**  high_imid  low_both  low_glypho  low_imid  high  low_glypho  low_imid  low_both  **high_glypho**  high_imid  **high_both**  high | **1.26 (0.17)**  **1.05 (0.17)**  0.06 (0.17)  0.26 (0.17)  0.44 (0.19)  0.15 (0.17)  0.10 (0.10)  4.8  1.4  1.0  1.0  **3.9**  1.0  **3.9**  1.0  3.0 | **7.50**  **6.25**  0.38  1.52  2.37  0.87  0.99  2.92  5.43  1.05  0.57  **19.49**  0.07  **20.66**  0.01  0.14 | **<0.001***  **<0.001***  0.707  0.134  0.021  0.389  0.328  0.021  0.007  0.309  0.453  **<0.001***  0.793  **<0.001***  0.941  0.24 |
| log_10_(*Flavobacterium*) | 89 | 0.48 | **treatment**  nutrient  ti(day)  **ti(day, by=treatment)**  ti(day, by=nutrient)  s(day, Pond, bs='fs') | **high_both**  **high_glypho**  high_imid  low_both  low_glypho  low_imid  high  low_glypho  low_imid  low_both  high_glypho  high_imid  **high_both**  high | **1.5 (0.25)**  **1.0 (0.25)**  0.24 (0.25)  0.46 (0.25)  0.51 (0.28)  0.16 (0.15)  -0.06 (0.15)  4.7  1.8  1.0  1.9  2.1  1.0  **1.0**  2.0  0.2 | **5.96**  **3.98**  0.95  1.82  1.80  0.64  -0.40  2.78  1.17  2.24  1.01  3.63  0.27  **17.35**  0.64  0.01 | **<0.001***  **<0.001***  0.348  0.074  0.076  0.522  0.687  0.042  0.288  0.139  0.325  0.031  0.606  **<0.001***  0.442  0.421 |
| log_10_(*Azospirillum*) | 89 | 0.78 | **treatment**  nutrient  ti(day)  **ti(day, by=treatment)**  ti(day, by=nutrient)  s(day, Pond, bs='fs') | **high_both**  **high_glypho**  high_imid  low_both  low_glypho  low_imid  high  low_glypho  low_imid  low_both  **high_glypho**  high_imid  **high_both**  high | **1.34 (0.17)**  **1.02 (0.17)**  0.09 (0.17)  0.31 (0.17)  0.18 (0.18)  0.18 (0.17)  -0.24 (0.10)  2.2  1.0  1.0  1.6  **2.6**  1.0  **3.3**  2.2  7.8 | **7.82**  **5.99**  0.50  1.83  0.98  1.06  -2.47  2.11  0.01  0.00  0.76  **5.41**  0.09  **6.27**  1.40  0.60 | **<0.001***  **<0.001***  0.62  0.07  0.33  0.29  0.02  0.137  0.920  0.988  0.480  **0.002***  0.770  **0.001***  0.192  0.015 |

^†^ When factor is absent it means the respective predictor variable is continuous (“day”) or a random effect (“pond”).

**Supplementary figures captions**

**Fig. S1** Experimental gradient established for (A) glyphosate and (B) imidacloprid concentrations between two application pulses (at days 6 and 34) and (C) the correlation between target and measured concentrations at each pulse. The top row of figure C shows results for glyphosate, and the bottom two rows for imidacloprid, after pulse 1 (left column) and pulse 2 (right column) respectively.

**Fig. S2** PRC plots show no effect of experimental treatments on community metabolic profiles when considering (A) the 31 different compounds individually (F=32.6 *p*=0.69) or (B) grouped according to functional guilds (F=79.2 *p*=0.86). The PRC axis shown in A explains 13.4% of total variance and in B 43.1%.

**Fig. S3** PCoA ordinations based on (A, B) weighted UniFrac distance or (B, D) Jensen-Shannon divergence exploring different normalization approaches: (A, C) calculation of reads relative abundance and (B, D) rarefying to a threshold of 10,000 reads per sample. Each sampling day is plotted separately to facilitate visualization of treatment effects on community composition.

**Fig. S4** PRCs show how high glyphosate treatments affected community composition at (A) phylum and (B) ASV levels. Low- and high-nutrient treatments show the same pattern, and, for this reason, they were grouped in Fig. 4, to facilitate data visualization. The finest level of taxonomic assignment based on FreshTrain and GreenGenes database is shown following ASV names in panel B. Only taxa with weights higher than 0.2 are shown in A and higher than 0.095 are shown in B.

**Fig. S5** PRCs show how high glyphosate treatments (15 mg/L) affected community composition at different taxonomic levels: (A) class, (B) order, (C) family/lineage, (D) genus/clade. Taxonomic assignment based on FreshTrain and GreenGenes databases. Low and high nutrient treatments were grouped as they follow the same pattern. Only taxa with weights higher than 0.2 are shown.

**Fig. S6** Summed effects of GAMMs on abundance of three genera most positively affected by the glyphosate treatments: (A) *Agrobacterium*, (B) *Flavobacterium* and (C) *Azospirillum*. Shades indicate a confidence interval of 95%. Abundance of each genus is the estimated absolute abundance of all ASVs assigned to *Agrobacterium*, *Flavobacterium* or *Azospirillum* after normalization by rarefying each sample to 10,000 reads without replacement.

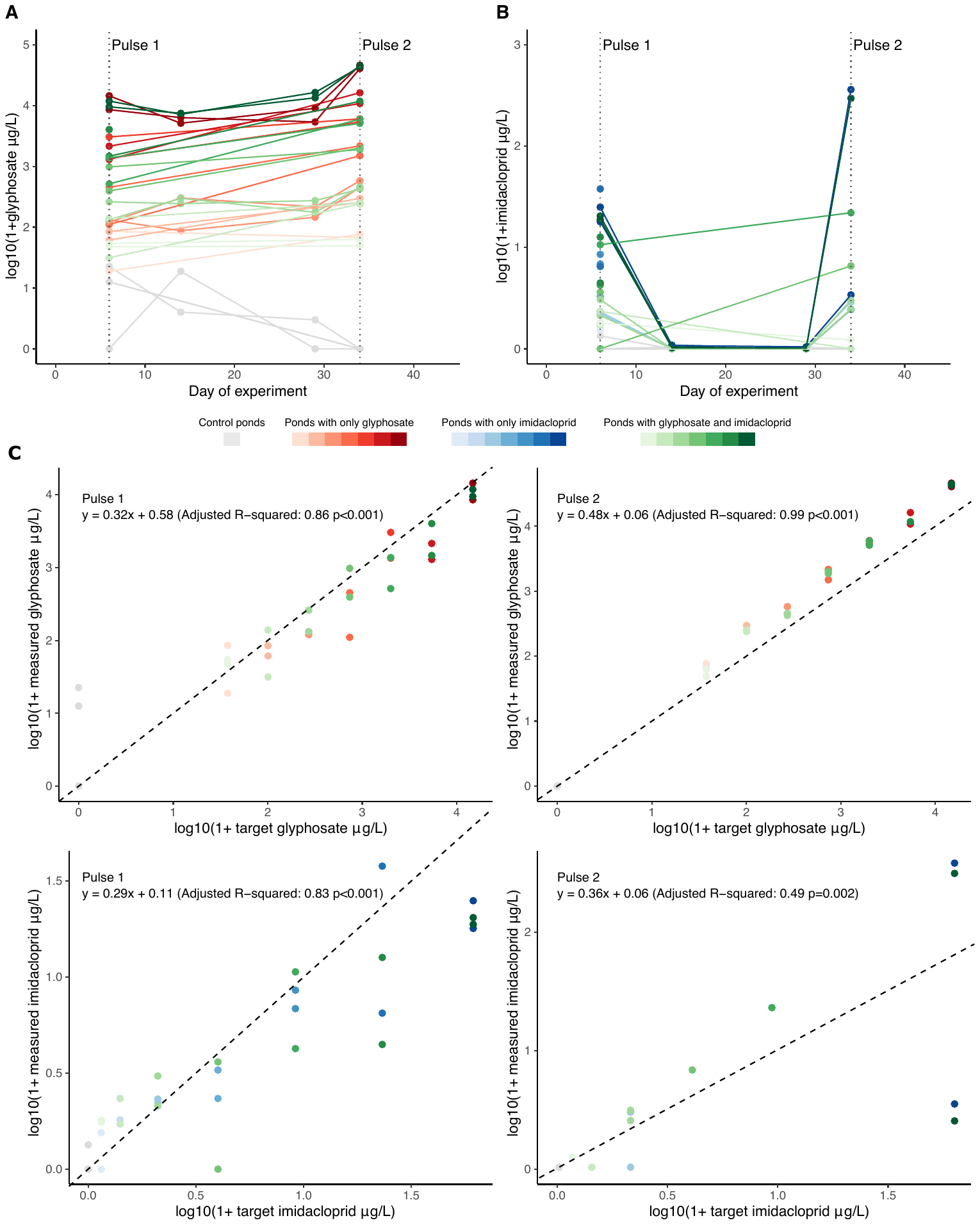

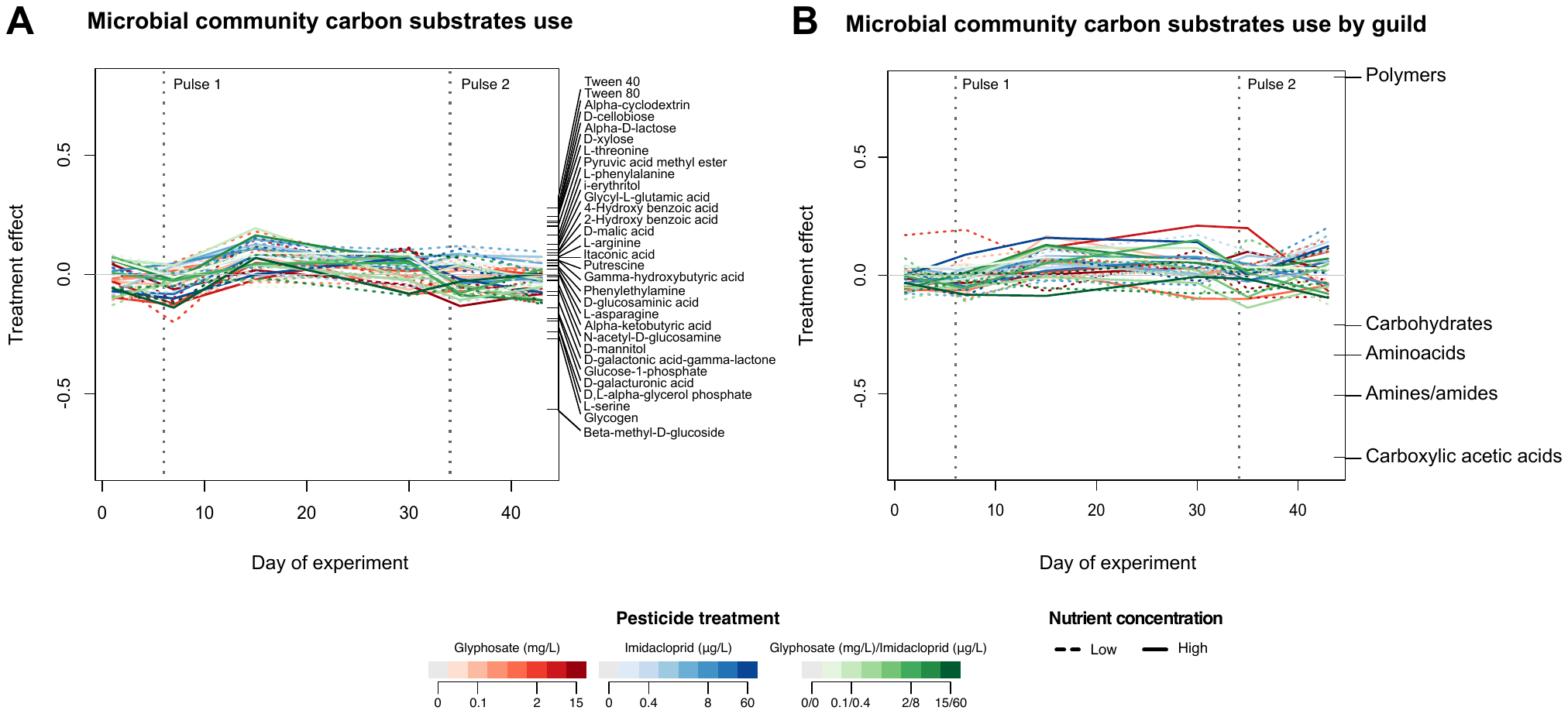

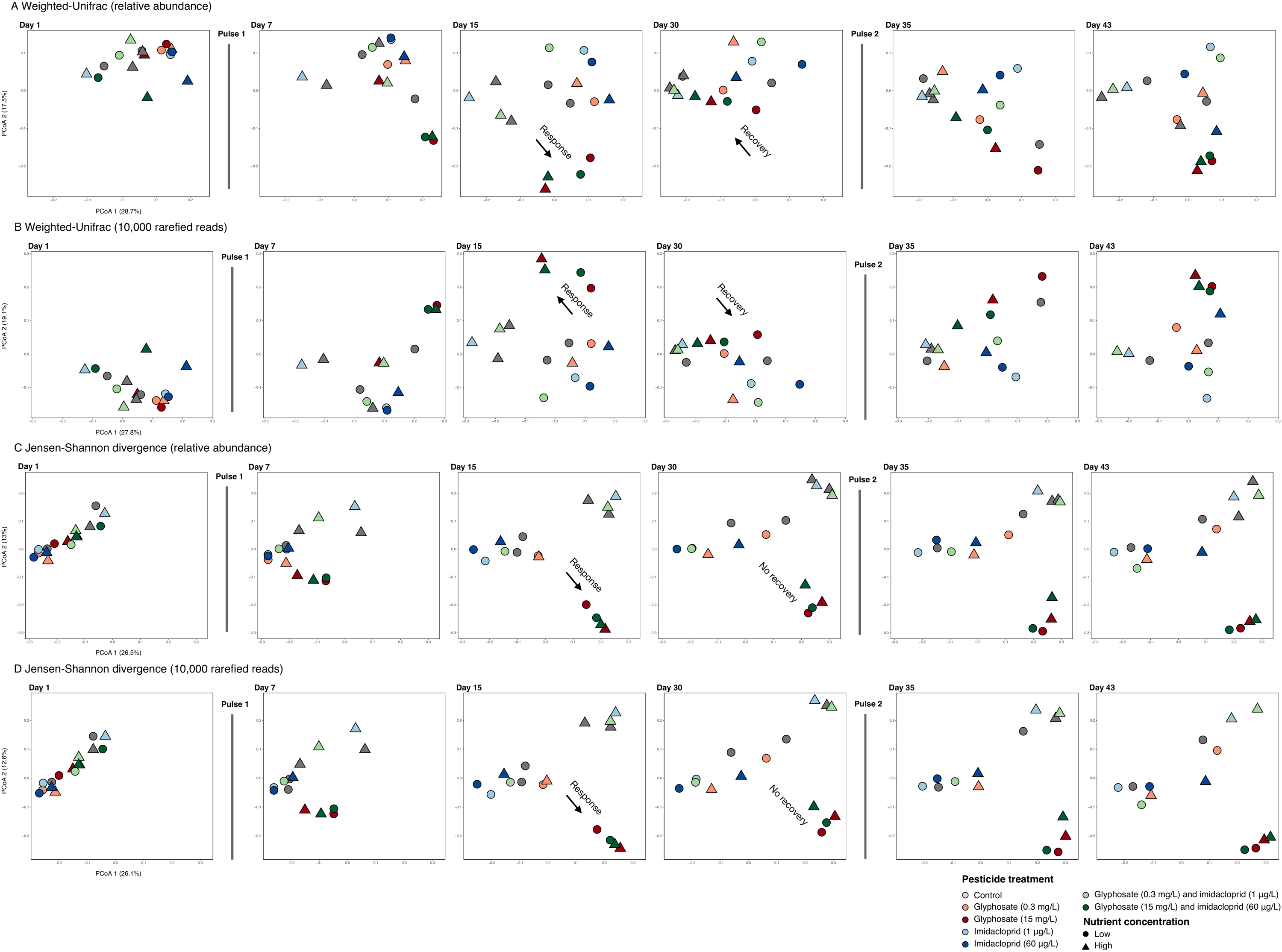

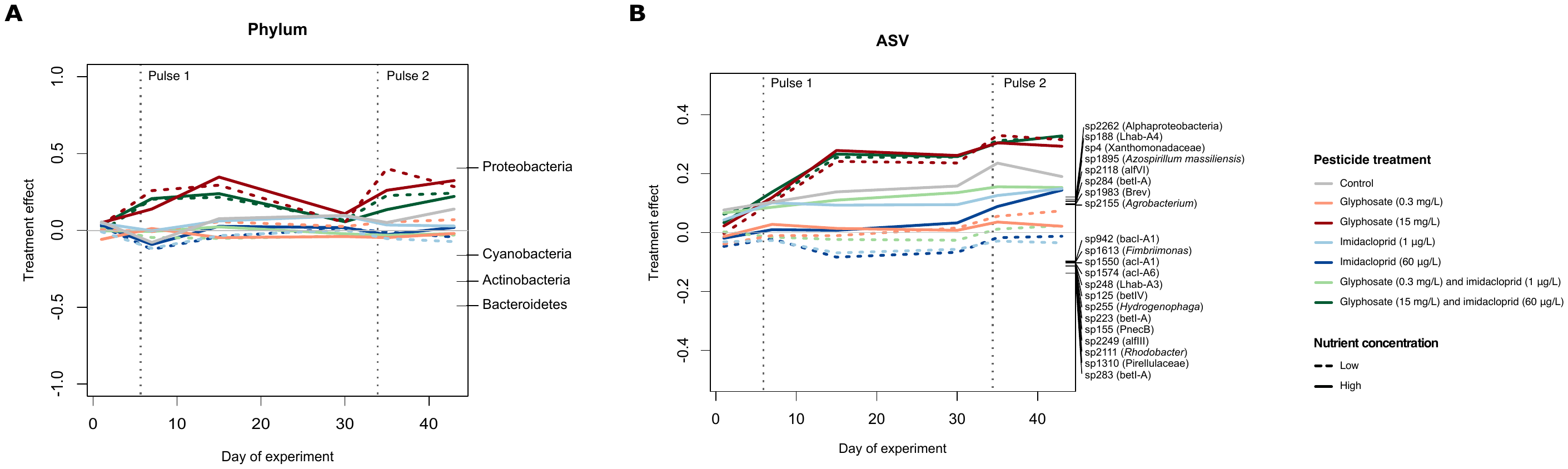

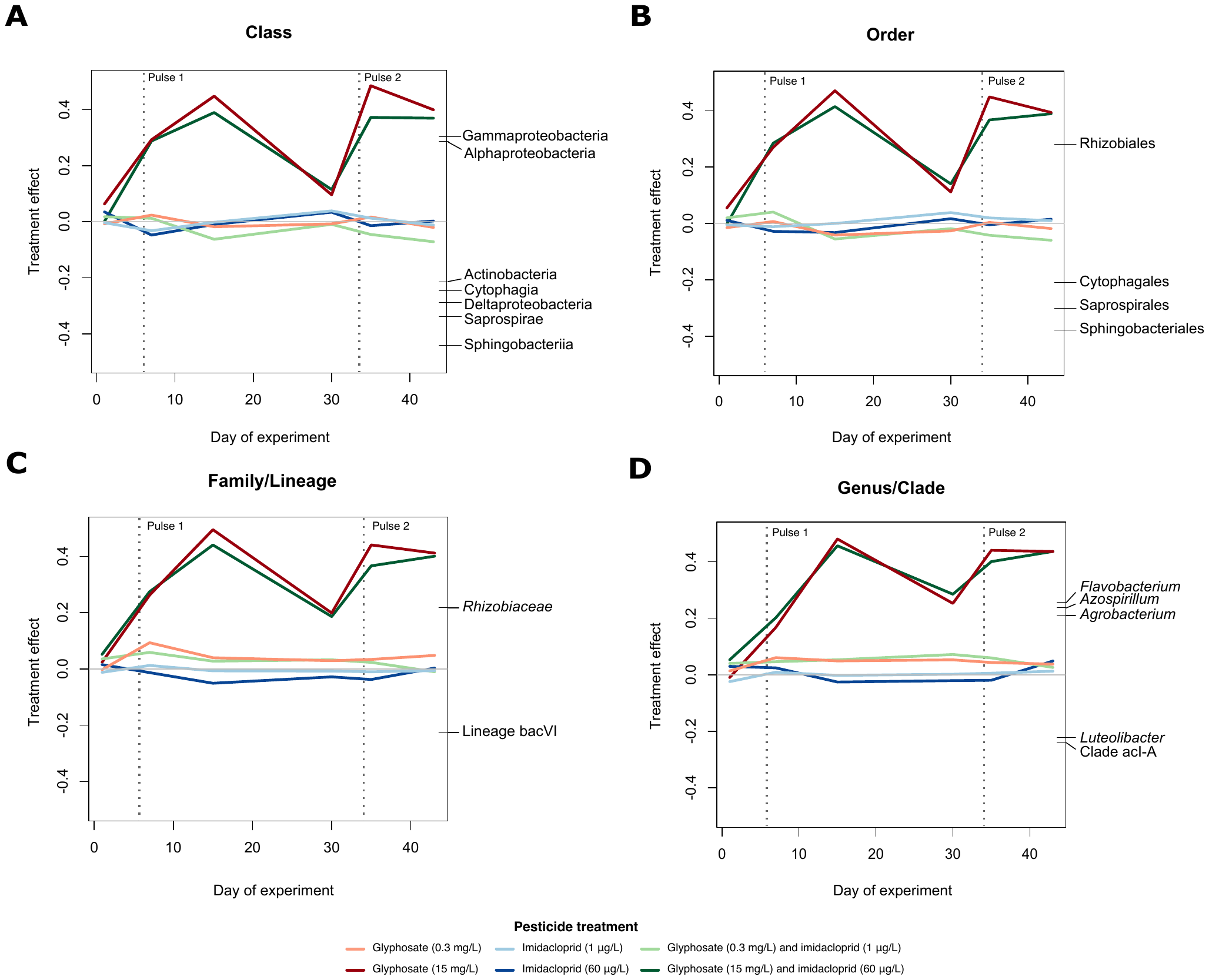

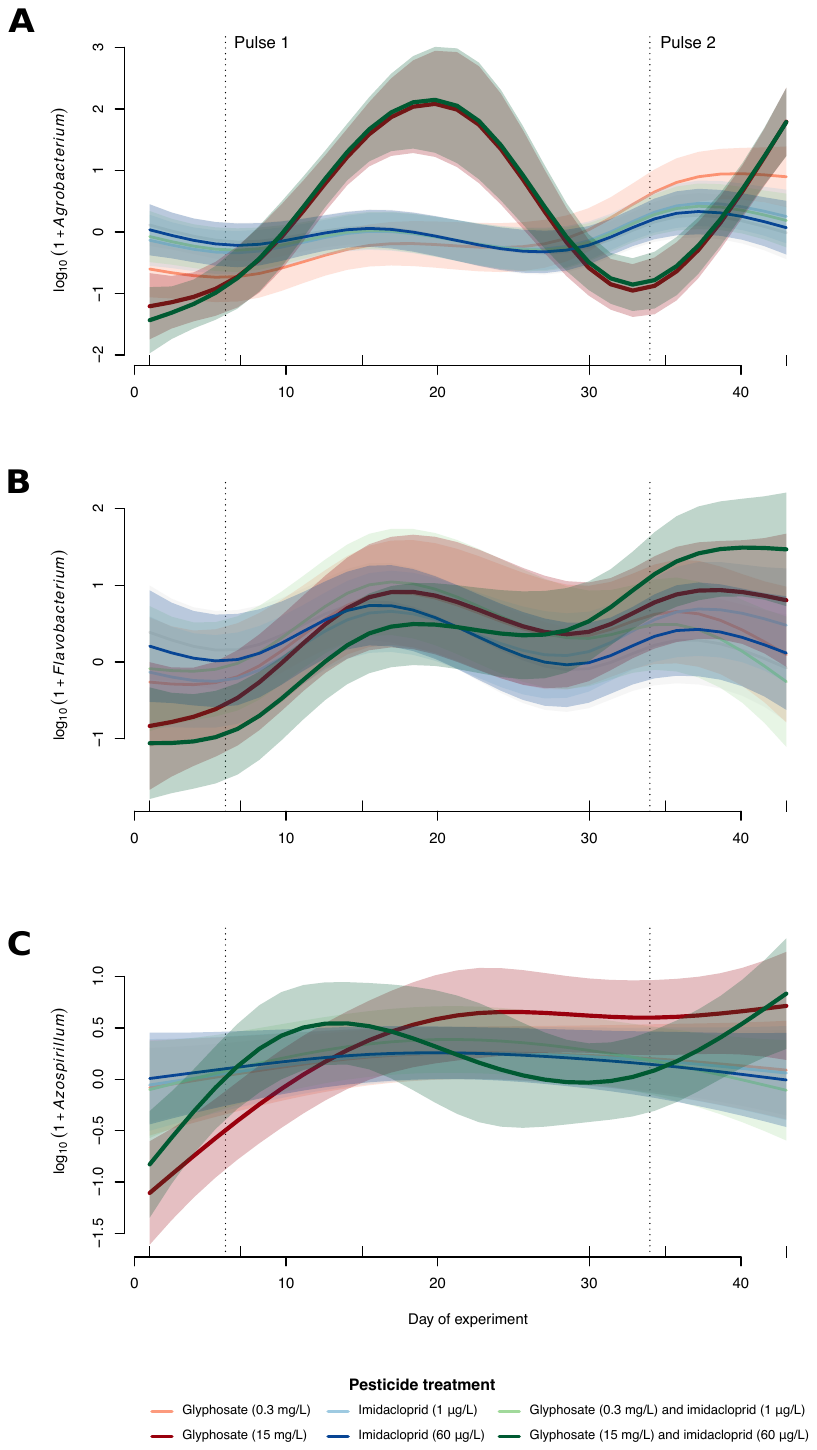
